# Oncogene-like addiction to aneuploidy in human cancers

**DOI:** 10.1101/2023.01.09.523344

**Authors:** Vishruth Girish, Asad A. Lakhani, Christine M. Scaduto, Sarah L. Thompson, Leanne M. Brown, Ryan A. Hagenson, Erin L. Sausville, Brianna E. Mendelson, Devon A. Lukow, Monet Lou Yuan, Pranav K. Kandikuppa, Eric C. Stevens, Sophia N. Lee, Barbora Salovska, Wenxue Li, Joan C. Smith, Alison M. Taylor, Robert A. Martienssen, Yansheng Liu, Ruping Sun, Jason M. Sheltzer

## Abstract

Most cancers exhibit aneuploidy, but its functional significance in tumor development is controversial. Here, we describe ReDACT (Restoring Disomy in Aneuploid cells using CRISPR Targeting), a set of chromosome engineering tools that allow us to eliminate specific aneuploidies from cancer genomes. Using ReDACT, we created a panel of isogenic cells that have or lack common aneuploidies, and we demonstrate that trisomy of chromosome 1q is required for malignant growth in cancers harboring this alteration. Mechanistically, gaining chromosome 1q increases the expression of MDM4 and suppresses TP53 signaling, and we show that TP53 mutations are mutually-exclusive with 1q aneuploidy in human cancers. Thus, specific aneuploidies play essential roles in tumorigenesis, raising the possibility that targeting these “aneuploidy addictions” could represent a novel approach for cancer treatment.

## INTRODUCTION

Chromosome copy number changes, otherwise known as aneuploidy, are a ubiquitous feature of tumor genomes^1, 2^. While the pervasiveness of aneuploidy in cancer has been known for over a century^3, 4^, the role of aneuploidy in tumor development has remained controversial^5–8^. Chromosome gains have been proposed to serve as a mechanism for increasing the dosage of tumor-promoting genes that are found within altered regions^9, 10^. However, proof of this hypothesis is lacking, and it has alternately been suggested that aneuploidy could arise as a result of the loss of checkpoint control that frequently occurs in advanced malignances^11^. Indeed, individuals with Down syndrome, which is caused by the triplication of chromosome 21, have a significantly decreased risk of developing most solid cancers, suggesting that in certain cases aneuploidy may actually have tumor-suppressive properties^12, 13^.

Our ability to directly interrogate the role of aneuploidy in cancer has historically been limited by the experimental difficulties involved in manipulating entire chromosome arms. Over the past 40 years, cancer researchers have used the standard tools of molecular genetics, including gene over-expression, knockdown, and mutagenesis, to develop a deep understanding of many individual oncogenes and tumor suppressors^14, 15^. For instance, the biological functions of genes like KRAS and TP53 were elucidated in part by creating and analyzing isogenic cell lines that express or lack these genes^16, 17^. However, existing approaches for single-gene manipulations are insufficient to interrogate the chromosome-scale changes that commonly occur in tumors and that affect hundreds or thousands of genes simultaneously. The consequences of eliminating specific aneuploid chromosomes from human cancer cells have not previously been established.

Studies of individual cancer driver genes led to the discovery of a phenomenon called “oncogene addiction”, in which loss or inhibition of a single oncogene is sufficient to induce cancer regression^18^. For example, mutations in KRAS cause the development of pancreas cancer, and genetically ablating KRAS in a “KRAS-addicted” pancreas tumor blocks growth and triggers apoptosis^19^. The principle of oncogene addiction also underlies the efficacy of cancer targeted therapies: drugs that inhibit “addictions” like EGFR and BRAF can result in sustained clinical responses in tumors that are driven by these oncogenes^20–22^.

Previous cancer genome sequencing projects have revealed that the aneuploidy patterns observed in human tumors are non-random, and specific chromosome gain events occur significantly more often than expected by chance^1, 23–25^. We speculated that these recurrent aneuploidies could themselves represent a novel type of cancer “addiction”, analogous to the concept of oncogene addictions. Eliminating these “aneuploidy addictions” could similarly block cancer growth and suppress malignant phenotypes. To investigate this hypothesis, we developed a set of computational and functional techniques to explore the similarities between aneuploidy and oncogenes and to uncover the phenotypic consequences of eliminating recurrent aneuploid chromosomes from established cancers.

## RESULTS

### Specific chromosome gains recurrently occur early in cancer development

Somatic mutations in oncogenes typically arise early in tumor evolution, consistent with their recognized roles as drivers of cancer development^26^. We recently developed a novel computational approach to leverage multi-sample tumor sequencing data to determine the relative timing of somatic copy number gains in cancer evolution^27^. We applied this tool to investigate the timing of aneuploidy events in a cohort of breast cancer (BRCA) and melanoma (MEL) patients^28, 29^. We found that specific chromosome copy number changes are consistently observed early in tumor development (Fig. 1A-B). Notably, we found that chromosome 1q gains are recurrently the first copy number alteration that occurs in breast cancer evolution, and these gains are also among the first alterations in melanoma evolution. In general, we observed that common aneuploidies arose earlier in tumor development than less-common aneuploidies, in agreement with the assumption that early somatic alterations are likely to be fitness-driving events (Fig. 1C)^30^. However, the correlation between frequency and timing was not maintained across all chromosomes. For instance, in breast cancer, chromosome 8q gains and chromosome 1q gains occur with similar frequencies, but we found that 1q gains consistently arose earlier during tumor development than 8q gains. We conclude that, as previously observed with oncogenic point mutations, specific chromosome gains occur in a defined temporal order, and we speculate that aneuploidies that are consistently gained early during tumorigenesis may enhance cancer fitness.

**Figure 1.**
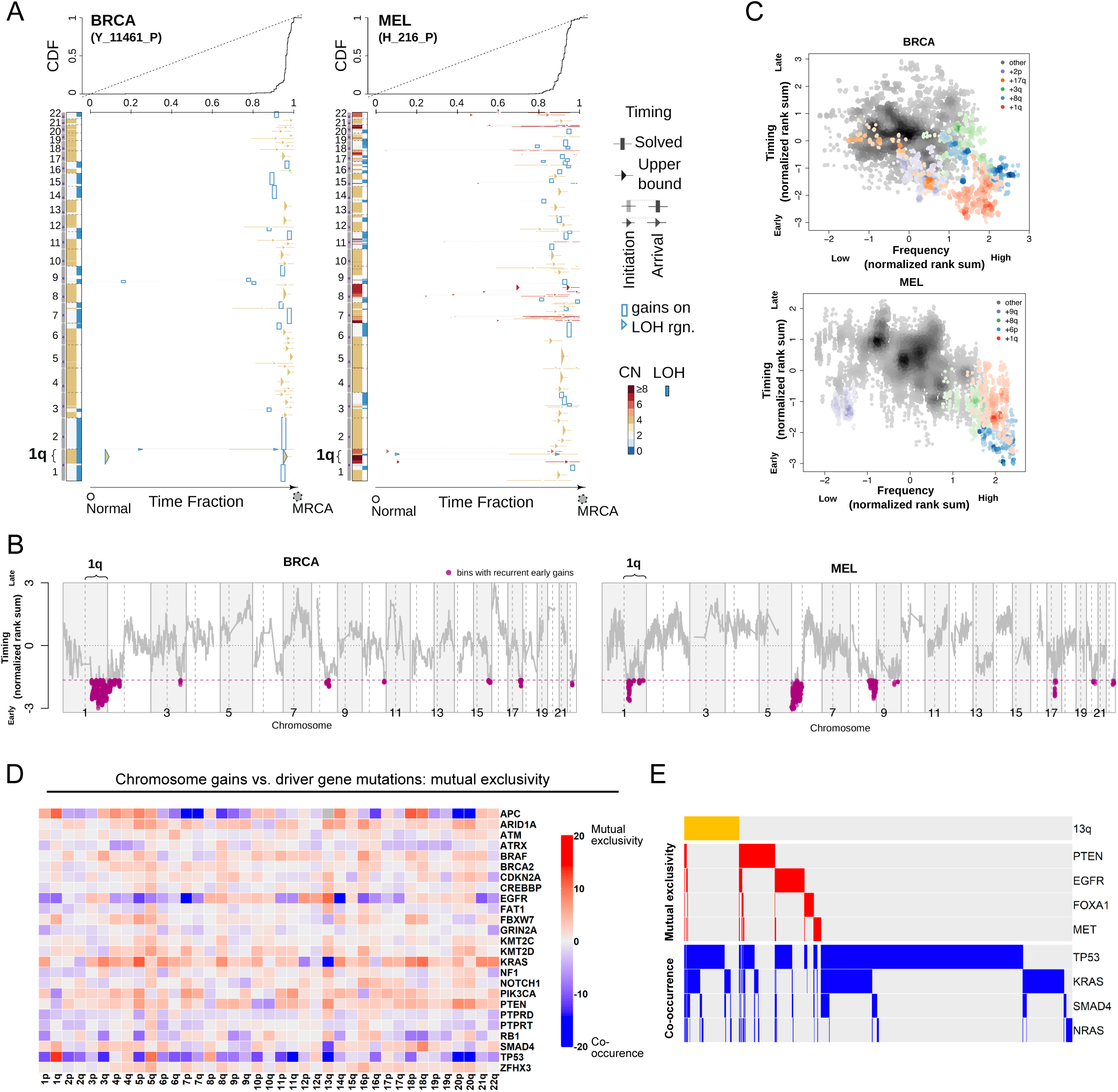
Specific chromosome gains arise early in tumor development and are mutually-exclusive with driver gene mutations. (A) The inferred timing of somatic copy number gains in the evolution of two tumors. A breast tumor is shown on the left and a melanoma on the right. Copy number (CN) states along the genome are shown on the left in each panel and color coded. The plot visualizes the time fraction of somatic evolution from germline to the most recent common ancestor (MRCA) of the patient tumor sample. For each copy number segment, the inferred timing is shown as a rectangle (exactly solved timing) or an arrow (upper bounds of timing when the timing solutions are not unique) with the same color-coding as its CN. The top panel shows the cumulative distribution (CDF) of the timing. Genome doubling (GD) can be observed as the punctuated gains occurring in a narrow time window and chromosome 1q gains appear to be extremely early and preceding GD in these two tumors. (B) Recurrent early gains of chromosome 1q in BRCA and MEL. For each tumor type, we converted the timing of gains into ranks for genomic bins within a patient and computed the rank sums across patients for each bin. The normalized rank sums for each genomic bin are shown for BRCA and MEL, respectively. The large negative values indicate recurrent early initiating gains. We used the normalized rank sums to test against the null hypothesis (no regions show recurrent early gain across patients). Bins from chromosome 1q reject this null for both tumor types (with 90% confidence level). (C) The timing of a gain compared to the frequency of its occurrence in BRCA and MEL. The points on the plots show the timing of gain of a genomic bin versus its frequency of copy number gain. Colors represent chromosomal arms, and color darkness indicates the density of points. Both the timings and frequencies are transformed into normalized rank sums (see Methods). (D) A pan-cancer analysis of mutual exclusivity between mutations in 25 commonly-mutated cancer genes and chromosome arm gain events. The complete results of this analysis are included in Table S1. (E) Mutual exclusivity and co-occurrence patterns between one representative chromosome gain (+13q, orange bars at the top), and point mutations in several different cancer driver genes.

### Specific chromosome gains are associated with altered mutational patterns and with cancer progression

In instances where two oncogenes converge to activate the same pathway, cancers frequently acquire mutations in either gene but not both^31, 32^. For example, melanomas can develop mutations in BRAF or NRAS that drive MAPK signaling, but co-mutation of both genes is redundant and rarely observed^33^. If chromosome gains play an oncogene-like role supporting cancer growth, then specific aneuploidies may also be expected to exhibit mutual exclusivity with individual oncogenic mutations. To investigate this possibility, we calculated patterns of mutual exclusivity between chromosome arm gains and mutations across 23,544 cancer patients^34^. We detected several hundred instances in which aneuploidies and mutations co-occur less often than expected by chance both within individual cancer types and in a pan-cancer analysis (Fig. 1D-E, S1, and Table S1). For instance, KRAS mutations are mutually exclusive with chromosome 18q gains in pancreatic cancer, while BRAF mutations are mutually exclusive with chromosome 20q gains in colorectal cancer (Fig. S1 and Table S1). These results are consistent with our hypothesis that specific chromosome gains can play an oncogene-like role in cancer, thereby making the acquisition of certain oncogenic mutations redundant in the presence of that aneuploidy.

High levels of aneuploidy are generally associated with poor cancer patient outcomes^35–37^. However, it is less clear whether specific copy number changes drive tumor progression, or whether the aneuploid state itself represents a universal risk factor. We calculated the association between patient outcome and copy number gains affecting every chromosome band across 10,884 patients and 33 cancer types from The Cancer Genome Atlas (TCGA). We discovered that certain copy number alterations were commonly prognostic across multiple cancer types, particularly gains affecting chromosome 1q (Fig. S2A-C and Table S2A). The strong association between 1q gains and disease progression was robust to the inclusion of multiple clinical variables, including patient age, sex, tumor stage, and tumor grade (Fig. S2D and Table S2B). 1q copy number correlated with hallmarks of aggressive disease in genetically-diverse cancer types, including with Gleason score in prostate adenocarcinoma and with thrombocytopenia in acute myeloid leukemia (Fig. S2E). We performed a similar analysis for cancer-associated mutations, and we found that the only gene for which mutations were prognostic in more than four cancer types is TP53 (Fig. S2A and S2D). These results illustrate that specific chromosome gain events, particularly involving regions of chromosome 1q, are robust pan-cancer markers for the risk of disease progression.

### Loss of trisomy 1q blocks malignant growth in human cancers

The computational analyses described above highlighted several similarities between chromosome copy number gains and driver mutations, raising the possibility that these aneuploidies could represent oncogene-like cancer addictions. The oncogene addiction paradigm was first established by developing genetic techniques to eliminate individual genes from established cancer cell lines^17, 18, 38^. In order to conduct comparable assays with aneuploidy, we created a set of approaches collectively called ReDACT (<u>Re</u>storing <u>D</u>isomy in <u>A</u>neuploid cells using <u>C</u>RISPR <u>T</u>argeting) (Fig. 2A). In the first approach, called ReDACT-NS (<u>N</u>egative <u>S</u>election), we integrate a single copy of herpesvirus thymidine kinase (HSV-TK) onto an aneuploid chromosome of interest. Then, the cells are transfected with a gRNA that cuts between the integrant and the centromere and treated with ganciclovir, which is toxic to cells that express HSV-TK^39^. Loss of the aneuploid chromosome harboring HSV-TK allows cells to survive ganciclovir selection. In the second approach, called ReDACT-TR (<u>T</u>elomere <u>R</u>eplacement), cells are co-transfected with a gRNA that cuts near the centromere of an aneuploid chromosome and a cassette encoding ∼100 repeats of the human telomere seed sequence^40^. CRISPR cleavage coupled with integration of the telomeric seed sequence leads to loss of an aneuploid chromosome arm and formation of a *de novo* telomere. In the third approach, called ReDACT-CO (<u>C</u>RISPR <u>O</u>nly), we took advantage of prior reports demonstrating that in rare circumstances CRISPR cleavage by itself is sufficient to trigger chromosome loss^41, 42^, and we transfected cells with a gRNA targeting an aneuploid chromosome arm without any other selection markers. We successfully applied all three approaches to create clones derived from human cell lines that had lost specific aneuploid chromosomes.

**Figure 2.**
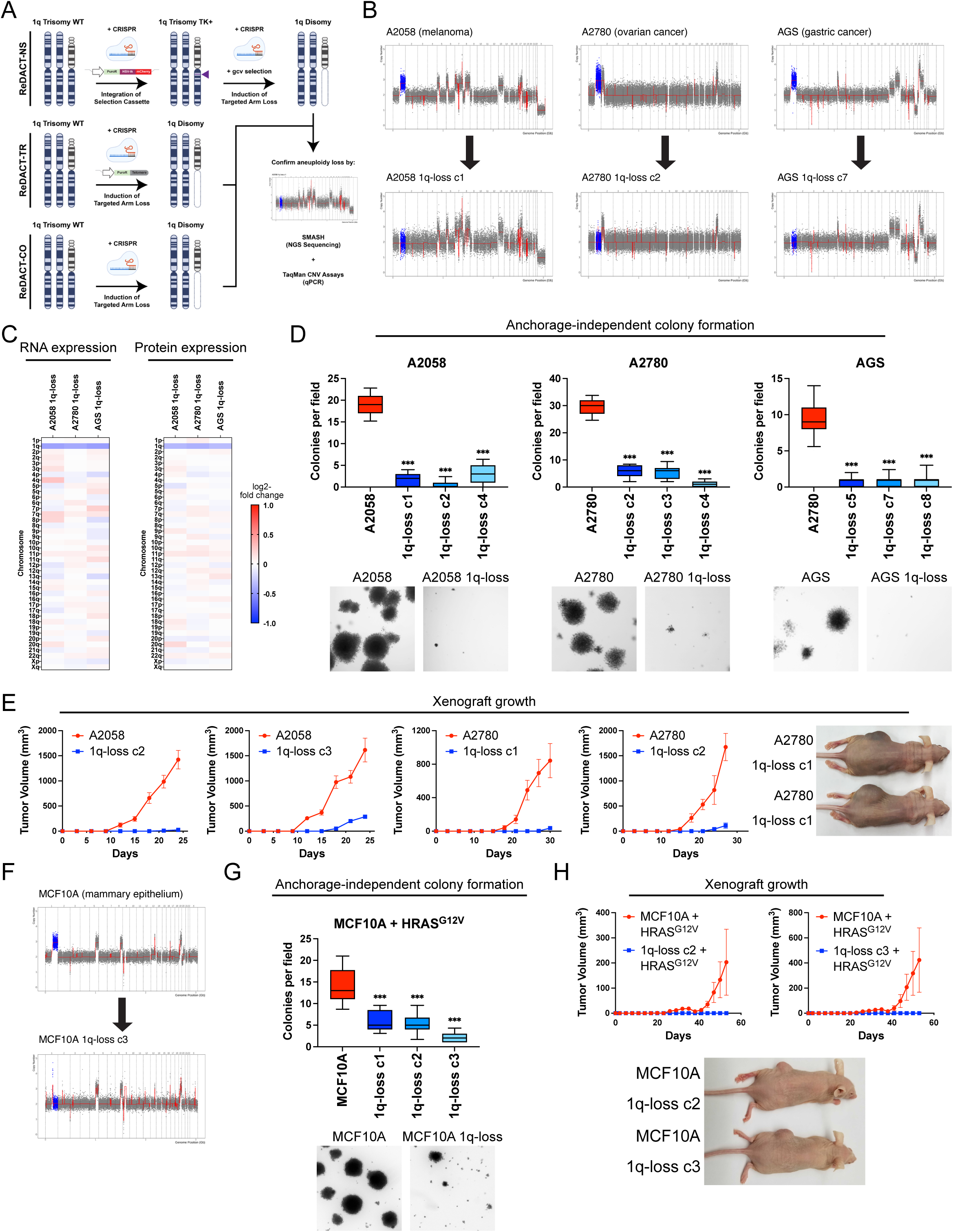
Phenotypic effects of losing chromosome 1q-aneuploidy. (A) Chromosomal engineering strategies for the targeted deletion of chromosome arms: (1) ReDACT-NS: using CRISPR-Cas9 homology-directed repair, we integrated a positive-negative selection cassette encoding a fluorescent reporter, a positive selection marker, and a negative selection marker (HSV thymidine kinase) at a centromere-proximal region on chromosome 1q. We induced arm loss by generating a dsDNA break centromere- proximal to the cassette with Cas9, and isolated clonal populations of cells that were ganciclovir-resistant. (2) ReDACT-TR: We induced arm loss by generating a dsDNA break at a centromere-proximal location with Cas9 while providing cells with an ectopic telomere seed sequence for repair. (3) ReDACT-CO: We induced arm loss by generating a dsDNA break at a centromere-proximal location with Cas9, and isolated clonal populations of cells. For all three approaches, we screened clonal populations of cells for targeted chromosome loss through TaqMan CNV assays and validated their karyotypes through SMASH sequencing. (B) Representative SMASH karyotypes of the 1q-disomic clones generated from the 1q-trisomic cancer cell lines A2780, AGS, and A2058. Chromosome 1q is highlighted in blue. A complete list of aneuploidy-loss clones is included in Table S3. (C) 1q-disomic clones display decreased RNA expression and protein expression of genes encoded on chromosome 1q. RNA expression data was obtained through bulk RNA-seq and represents the average expression of genes by chromosome arm across multiple 1q-disomic clones for each cell line. Protein expression data was obtained through mass spectrometry, and representative data from one 1q-disomic clone is shown for each cell line. Data are log2 transformed, normalized to the parental cell line, and adjusted so that the mean expression across all chromosomes is 0. (D) 1q-disomic clones exhibit decreased anchorage-independent growth. The micrographs display representative images of colony formation for 1q-trisomic and 1q-disomic clones. (E) 1q-disomic clones exhibit impaired xenograft growth *in vivo*. 1q-trisomic and 1q-disomic cells were injected contralaterally and subcutaneously into immunocompromised mice. The graphs display the mean ± SEM for each trial. Representative mice are shown on the right. (F) SMASH karyotype of a 1q-disomic clone generated from the mammary epithelial cell line MCF10A. Chromosome 1q is highlighted in blue. (G) 1q-disomic MCF10A clones transduced with HRAS^G12V^ exhibit decreased anchorage-independent growth relative to 1q-trisomic MCF10A cells. (H) 1q disomic MCF10A clones transduced with HRAS^G12V^ clones exhibit impaired xenograft growth *in vivo*. 1q- trisomic and 1q-disomic cells were injected contralaterally and subcutaneously into immunocompromised mice. The graphs display the mean ± SEM for each trial. Representative mice are shown below. For anchorage-independent growth assays in D and G, the boxplots represent the 25th, 50th, and 75th percentiles of colonies per field, while the whiskers represent the 10th and 90th percentiles. Unpaired t-test, n = 15 fields of view, data from representative trial. Representative images are shown below. **p < 0.005, ***p < 0.0005

We first focused on aneuploidies of chromosome 1q, as we found that 1q gains were an early event in multiple cancer types and were strongly associated with disease progression (Fig. 1 and S2). We targeted the 1q trisomy in the A2058 melanoma cell line, AGS gastric cancer cell line, and A2780 ovarian cancer cell line. We generated multiple independent derivatives of each line in which a single copy of chromosome 1q had been eliminated, thereby producing cell lines that were disomic rather than trisomic for chromosome 1q. We verified loss of the 1q trisomy and the absence of any other chromosome copy number changes using SMASH-Seq, a sequencing-based approach to determine DNA copy number^43^, and by G-banding analysis of metaphase spreads (Fig. 2B, S3-S4, and Table S3). Loss of the 1q trisomy decreased the expression of genes encoded on chromosome 1q by an average of 26% at the RNA level and 21% at the protein level (Fig. 2C). These results suggest that chromosome loss causes a substantial downregulation of genes encoded on an aneuploid chromosome, though these effects can be buffered to some extent by cellular dosage compensation^44, 45^.

Next, we tested whether losing the 1q trisomy affects malignant growth in cancer cells. Toward that end, we quantified anchorage-independent colony formation, an *in vitro* proxy for malignant potential^46, 47^, in the 1q- trisomic and 1q-disomic cells. While 1q-trisomic A2058, A2780, and AGS cells displayed robust colony formation, multiple independent 1q-disomic clones derived from each cell line exhibited minimal anchorage-independent growth (Fig. 2D). We then performed contralateral subcutaneous injections with each cell line to test whether aneuploidy-loss affected tumor growth *in vivo*. Consistent with our colony formation assays, we observed that 1q-trisomic A2058 and A2780 cells rapidly formed large tumors, while 1q-disomic cells displayed minimal tumor growth (Fig. 2E). At the end of these assays, the trisomic cells had formed tumors that were on average 25-fold larger than the tumors formed by the 1q-disomic cells. For the AGS cancer cell line, neither the trisomic nor the disomic cells formed tumors following subcutaneous injection (Fig. S5). Finally, we performed proliferation assays to measure the doubling time of the 1q-trisomic and the 1q-disomic cells in culture (Fig. S6A-C). The aneuploidy-loss clones divided more slowly *in vitro* compared to the 1q-trisomic cells, though the difference in doubling time (∼35%) was substantially less than the differences observed in the soft agar and xenograft assays. In total, these results suggest that multiple human cancer cell lines are dependent on the presence of a third copy of chromosome 1q to support malignant growth, and elimination of this aneuploid chromosome compromises their tumorigenic potential. Furthermore, we note that this phenotypic pattern, in which aneuploidy loss causes a moderate effect on *in vitro* doubling but a severe effect on anchorage-independent growth and xenograft formation, resembles the previously-reported consequences of eliminating *bona fide* oncogene addictions^17, 48^.

### Loss of trisomy 1q prevents malignant transformation

We discovered that 1q gains were commonly the first copy number alteration to occur during breast tumor development (Fig. 1A-C). We therefore hypothesized that, in addition to being required for cancer growth, aneuploidy of chromosome 1q may directly promote cellular transformation. To test this, we performed chromosome engineering in MCF10A, an immortal but non-tumorigenic mammary epithelial cell line^49^. SMASH- Seq revealed that this cell line harbors a trisomy of chromosome 1q, and we successfully applied ReDACT-CO and ReDACT-TR to generate derivatives of MCF10A with two copies rather than three copies of 1q (Fig. 2F, S3D, and Table S3). We then attempted to transform the 1q-trisomic and 1q-disomic cells by transducing them with a retrovirus encoding the HRAS^G12V^ oncogene. HRAS^G12V^ expression was sufficient to transform trisomic MCF10A, as these cells were able to form colonies in soft agar and grow as xenografts in nude mice (Fig. 2G- H). In contrast, 1q-disomic MCF10A clones expressing HRAS^G12V^ exhibited impaired colony formation and were unable to produce tumors *in vivo*, demonstrating that loss of the trisomic chromosome prevented cellular transformation. These results are consistent with our finding that 1q gains are an early event during breast cancer development and demonstrate that specific aneuploidies can cooperate with oncogenes to transform non-malignant cells.

### Robust anchorage-independent growth in human cancer cell lines subjected to CRISPR cutting and ganciclovir selection

In order to confirm that our findings were a specific consequence of aneuploidy loss, we generated and tested a series of control clones subjected to various CRISPR manipulations that did not induce loss of the 1q trisomy (Fig. S7 and S8A). These control clones included:

1) Cell lines harboring a CRISPR-mediated integration of the HSV-TK cassette that were not subjected to selection for 1q-loss
2) Cell lines in which the HSV-TK cassette was deleted by transfecting cells with two gRNAs targeting immediately upstream and downstream of the integrant coupled with ganciclovir selection, which resulted in a segmental deletion of the cassette while leaving the rest of 1q unaffected
3) Cell lines subjected to CRISPR-mediated cutting with a 1q-targeting gRNA, in which the lesion was repaired without inducing chromosome loss
4) Cell lines subjected to CRISPR-mediated cutting with a gRNA targeting the non-coding Rosa26 locus^50^
5) Cell lines in which dual CRISPR guides were used to generate segmental deletions on chromosome 1q of genes encoding non-expressed olfactory receptors
6) Cell lines in which CRISPR was used to delete a terminal segment on chromosome 1q, eliminating the telomere and decreasing the copy number of 26 out of 968 protein-coding genes on the chromosome

We performed SMASH-Seq on each control clone that we generated and we confirmed that each clone retained an extra copy of chromosome 1q (Fig. S7). We then measured the effects of these manipulations on anchorage-independent growth. We found that every control clone maintained the ability to form colonies in soft agar, with some variability between independent clones. Across the three cancer cell lines and 28 different control clones, we observed that the 1q-disomic clones exhibited significantly worse anchorage-independent growth than every control clone that we generated (Fig. S8B-E). These results indicate that the deficiencies in malignant growth exhibited by the 1q-disomic clones are not a result of our use of CRISPR or ganciclovir selection (discussed in more detail below and in Supplemental Text 1).

### Eliminating different cancer aneuploidies produces unique phenotypic consequences

We investigated two alternate possibilities to explain the loss of malignant potential that we observed following elimination of the 1q trisomy. First, we considered the possibility that the poor growth of the 1q-disomic clones was a consequence of some aspect of the ReDACT approach that was not fully captured by the control clones tested above. Second, while we initially focused on aneuploidy of chromosome 1q due to its occurrence early in cancer development and its association with poor prognosis (Fig. 1A-C and S2), we considered the possibility that the loss of any clonal aneuploid chromosome from a cancer cell line could have identical phenotypic effects. To further investigate the consequences of inducing aneuploidy-loss, we took advantage of the A2058 melanoma cell line, which harbors multiple trisomic chromosomes. We previously described the consequences of eliminating trisomy 1q from this cell line (Fig. 2); we subsequently used the same ReDACT techniques to generate derivatives of this cell line that had lost either the trisomy of chromosome 7p or the trisomy of chromosome 8q. SMASH-Seq confirmed the desired aneuploidy-loss events without other karyotypic changes (Fig. 3A, S9A-B, and Table S3).

**Figure 3.**
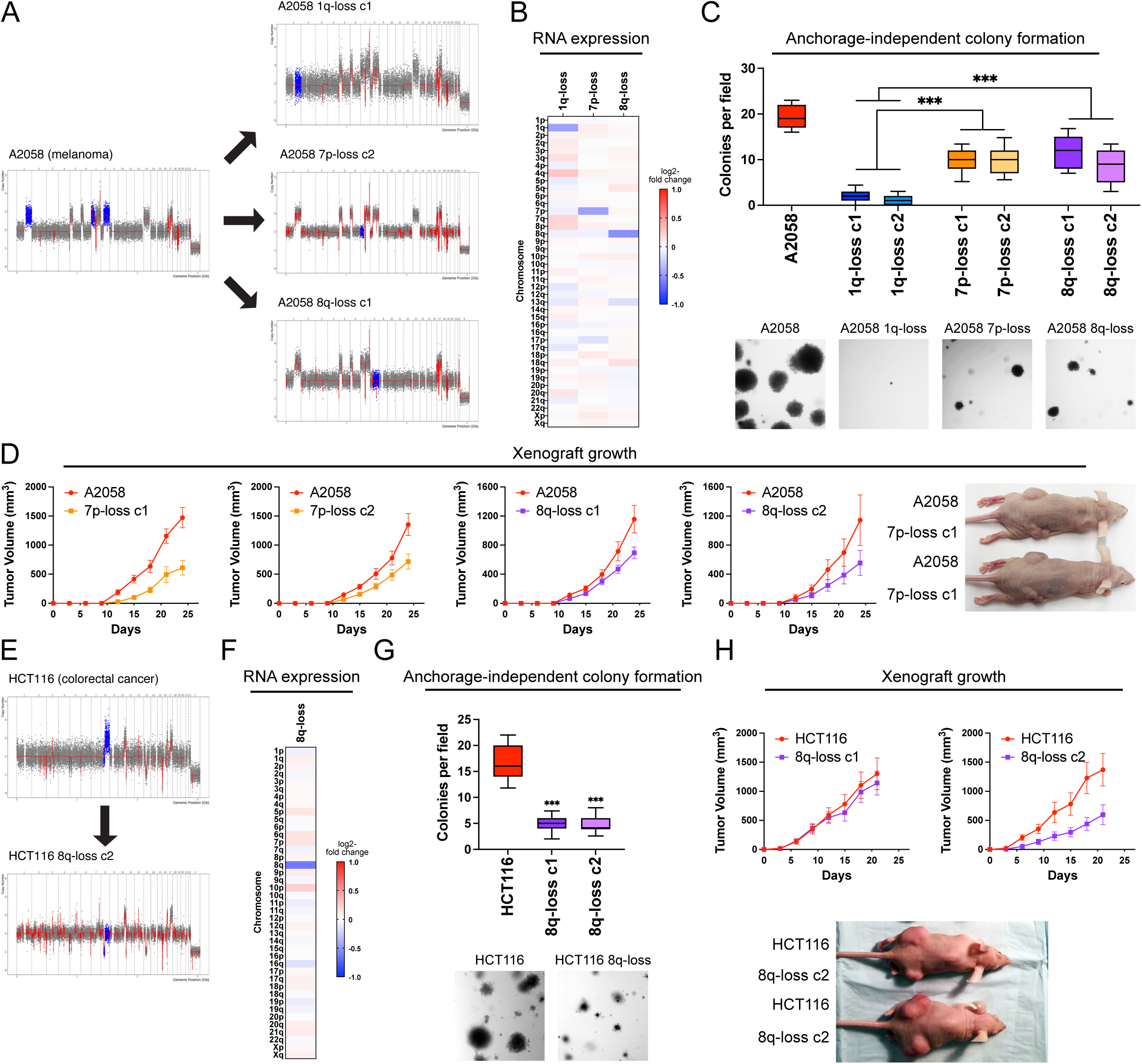
Variable degrees of addiction to aneuploidy of chromosome 1q, 7p, and 8q. (A) Representative SMASH karyotypes of the 1q-disomic, 7p-disomic, and 8q-disomic clones generated from the melanoma cell line A2058. Trisomy of chromosomes 1q, 7p, and 8q are highlighted in blue in the parental cell line on the left, and the respective targeted chromosome loss is highlighted in blue in the derived clones on the right. A complete list of aneuploidy-loss clones is included in Table S3. (B) 1q-disomic, 7p-disomic, and 8q-disomic clones in A2058 exhibit decreased RNA expression of genes encoded on the targeted chromosome. RNA expression data was obtained through bulk RNA-seq and represents the average expression of genes by chromosome arm across multiple aneuploidy-loss clones for each targeted chromosome. Data are log2 transformed, normalized to the parental cell line, and adjusted so that the mean expression across all chromosomes is 0. (C) 7p-disomic and 8q-disomic clones in A2058 exhibit a milder deficit in anchorage-independent growth as compared to 1q-disomic clones. The micrographs display representative images of colony formation for the indicated cell lines. (D) 7p-disomic and 8q-disomic clones in A2058 exhibit a moderate defect in xenograft growth. Wild-type (7p- trisomic and 8q-trisomic) cells and either 7p-disomic or 8q-disomic cells were injected contralaterally and subcutaneously into immunocompromised mice. The graphs display the mean ± SEM for each trial. Representative mice are shown on the right. (E) SMASH karyotype of an 8q-disomic clone generated from the colorectal cancer cell line HCT116. Chromosome 8q is highlighted in blue. (F) 8q-disomic clones in HCT116 exhibit decreased RNA expression of genes encoded on chromosome 8q. RNA expression data was obtained through bulk RNA-seq and represents the average expression of genes by chromosome arm across multiple aneuploidy-loss clones for each cell line. Data are log2 transformed, normalized to the parental cell line, and adjusted so that the mean expression across all chromosomes is 0. (G) 8q-disomic clones in HCT116 exhibit decreased anchorage-independent growth. The micrographs display representative images of colony formation for the indicated cell lines. (H) 8q-disomic clones in HCT116 exhibit variable xenograft growth. 8q-trisomic and 8q-disomic cells were injected contralaterally and subcutaneously into immunocompromised mice. The graphs display the mean ± SEM for each trial. Representative mice are shown below the graphs. For anchorage-independent growth assays in C and G, boxes represent the 25th, 50th, and 75th percentiles of colonies per field, while the whiskers represent the 10th and 90th percentiles. Unpaired t-test, n = 15 fields of view, data from representative trial. Representative images are shown below. ***p < 0.0005

As expected, loss of either trisomy 7p or trisomy 8q resulted in a significant decrease in the expression of genes encoded on the affected chromosomes (Fig. 3B). Next, we tested anchorage-independent growth in these aneuploidy-loss clones. If the poor growth of the 1q-disomic clones is a non-specific effect of ReDACT, then we would expect that all aneuploidy-loss clones would exhibit similar phenotypes. While 7p-disomic and 8q-disomic clones exhibited impaired anchorage-independent growth compared to a panel of control clones, we observed that this defect was not as severe as the defect observed among 1q-disomic clones (Fig. 3C and S10). Loss of trisomy-1q resulted in a 92% decrease in colony formation relative to the parental A2058 cells, compared to decreases of 49% and 47% for 7p-loss and 8q-loss, respectively. *In vitro* doubling times were also closer to wild-type levels for 7p-disomic and 8q-disomic cells compared to 1q-disomic cells (Fig. S6D). Finally, we measured tumor growth following subcutaneous injections of the 7p-disomic and the 8q-disomic cells in nude mice. Loss of either the 7p or the 8q trisomy resulted in a moderate decrease in tumor growth (Fig. 3D). At the end of the assay, the wild-type tumors were on average two-fold larger than the tumors formed by either 7p-disomic or 8q- disomic cells, compared to a 30-fold difference between A2058 wild-type and 1q-disomic tumors (Fig. 2E). In total, these results indicate that A2058 melanoma cells exhibit a greater degree of “addiction” to the 1q-trisomy compared to the trisomies of chromosome 7p or 8q.

To explore the consequences of losing chromosome 8q aneuploidy in a distinct cancer type, we eliminated the 8q-trisomy from the colorectal cancer cell line HCT116 (Fig. 3E-F, S9C and Table S3). Consistent with our observations in A2058, loss of the 8q trisomy significantly decreased but did not fully prevent anchorage-independent growth in HCT116 (Fig. 3G). We then tested xenograft formation in the HCT116 8q-disomic cells, and we observed that one 8q-loss clone exhibited a moderate defect in tumor growth while a second clone was able to form tumors at levels comparable to the trisomic parental line (Fig. 3H). These results demonstrate that eliminating aneuploid chromosomes has variable effects, depending on the identity of the chromosome and the genetic background of the cancer.

### Karyotype evolution and 1q trisomy restoration after aneuploidy loss

A hallmark of oncogene addictions is that loss or inhibition of a driver oncogene results in strong and rapid selection to re-establish oncogenic signaling^51^. For instance, when EGFR-driven lung cancers are treated with an EGFR inhibitor, these cancers evolve to acquire specific mutations, including EGFR^T790M^ (which restores EGFR activity) and KRAS^G13D^ (which activates a parallel oncogenic pathway)^52^. We sought to investigate whether elimination of an “aneuploidy addiction” would also result in evolutionary pressure to restore the lost aneuploidy. We injected 1q-disomic A2058 cells into nude mice and then determined the copy number of chromosome 1q in the resulting xenografts using qPCR (Fig. 4A). We discovered that 65 out of 82 1q-disomic xenografts re-acquired an extra copy of chromosome 1q, demonstrating strong selective pressure to regain the initial 1q aneuploidy. We subjected 20 of these post-xenograft clones to SMASH-Seq, and we found that chromosome 1q-regain was the only detectable chromosome-scale copy number change (Fig. 4B and S11A). No gross karyotypic changes were observed when the parental 1q-trisomic cells were grown as xenografts (Fig. 4B and S11B). If loss of the chromosome 1q trisomy compromises malignant potential, then we would expect that regaining 1q aneuploidy would restore cell fitness. Consistent with this hypothesis, we found that cells that had re-acquired the 1q trisomy exhibited increased clonogenicity compared to the 1q-disomic cells when grown under anchorage-independent conditions (Fig. 4C).

**Figure 4.**
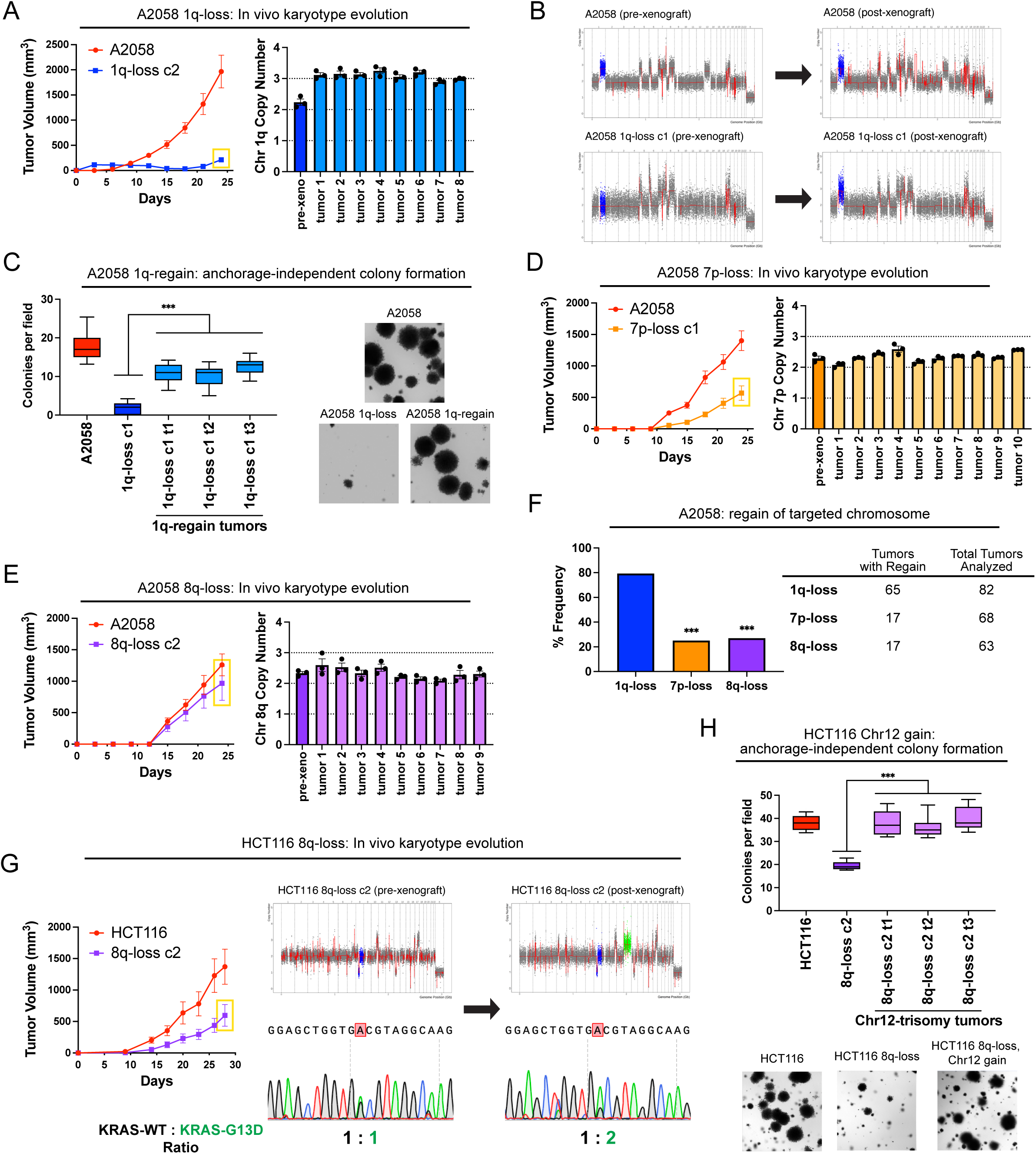
Cancers rapidly recover chromosome 1q aneuploidy. (A) A2058 1q-disomic cells frequently evolve to recover a third copy of chromosome 1q during xenograft growth. (B) Representative SMASH karyotypes of A2058 wildtype and 1q-disomic tumors. The initial karyotypes for these lines prior to the xenograft assay are shown on the left, and karyotypes of tumors following the xenograft assay are shown on the right. Chromosome 1q is highlighted in blue. (C) 1q-disomic clones that have evolved to regain 1q-trisomy following xenograft growth exhibit increased anchorage-independent growth relative to the pre-xenograft 1q-disomic parental cells. (D) Variable evolution of 7p-disomic cells to recover a third copy of chromosome 7p during xenograft growth. (E) Variable evolution of 8q-disomic cells to recover a third copy of chromosome 8q during xenograft growth. (F) Regain of trisomy 1q occurs more frequently than regain of trisomy 7p or trisomy 8q. Tumors were classified as exhibiting regain if the mean copy number of the targeted chromosome was ≥ 2.5, as determined through TaqMan copy number assays. n = 213 tumors, chi-squared test. (G) HCT116 8q-disomic clones evolve to gain a copy of chromosome 12 during xenograft assays, resulting in the acquisition of an extra copy of the KRAS^G13D^ allele. Cell lines were rederived from tumors harvested at the endpoint of xenograft assays, and subjected to SMASH karyotyping and Sanger sequencing of KRAS. The xenograft growth curve is shown on the left, and representative SMASH karyotype profiles and Sanger sequencing chromatograms pre- and post-xenograft are shown on the right. Chromosome 8q is highlighted in blue and chromosome 12 is highlighted in green. (H) 8q-disomic clones that have evolved to acquire trisomy of chromosome 12 following xenograft growth exhibit increased anchorage-independent growth relative to the pre-xenograft 8q-disomic parental cells. For copy number profiling in A, D, and E, cell lines were rederived from tumors at the endpoint of the xenograft assays, and chromosome copy number was determined through TaqMan copy number assays. Mean ± SEM, n = 3 probes on targeted chromosome, data from representative trials are shown. The corresponding xenograft assays are shown on the left. For the anchorage-independent growth assays in C and H, the boxes represent the 25th, 50th, and 75th percentiles of colonies per field, while the whiskers represent the 10th and 90th percentiles. Unpaired t-test, n = 15 fields of view, data from representative trial. Representative images are shown on the right. ***p < 0.0005

Next, we assessed karyotype evolution following *in vivo* growth of A2058 7p-disomic and 8q-disomic clones. Interestingly, 17 out of 68 7p-disomic xenografts and 17 out of 63 8q-disomic xenografts were found to exhibit 7p- and 8q-trisomy regain, respectively (Fig. 4D-E). These rates of chromosome re-gain were significantly lower than the rates that we observed for chromosome 1q (P < .0001, chi-square test; Fig. 4F). These results suggest that there is moderate selective pressure to restore 7p and 8q trisomies and stronger selective pressure to restore 1q trisomy in A2058.

We then sought to determine whether we could observe evolutionary pressure to restore chromosome 1q aneuploidy when 1q-disomic cells were grown *in vitro*. Toward that end, we passaged A2058, A2780, and AGS 1q-trisomic and 1q-disomic cancer cell lines for thirty days in culture and then we assessed their karyotypes. Similar to our *in vivo* results, we uncovered multiple instances in which 1q-disomic cells independently regained an extra copy of chromosome 1q over the course of the assay (Fig. S12).

Finally, we assessed karyotype evolution in xenografts produced by 8q-disomic HCT116 cells (Fig. 4G). Interestingly, we found that 0 out of 13 tumors regained the trisomy of chromosome 8q, but 7 out of 13 tumors gained a *de novo* trisomy of chromosome 12. HCT116 cells are driven by a heterozygous KRAS^G13D^ mutation^17^, and KRAS is encoded on chromosome 12. Sanger sequencing analysis revealed that every chromosome 12- trisomic tumor had amplified the copy of chromosome 12 harboring the mutant KRAS^G13D^ allele (Fig. 4G). Increasing dosage of mutant KRAS has previously been associated with enhanced tumor fitness^53, 54^, and we observed that these chromosome 8q-disomic/chromosome 12-trisomic cells exhibited superior anchorage-independent growth relative to the chromosome 8q-disomic/chromosome 12-disomic pre-xenograft population (Fig. 4H). In total, these results suggest that aneuploidy loss creates strong selective pressure for karyotype evolution, and the effects of aneuploidy loss can be suppressed *in cis* (by regaining the lost chromosome) or *in trans* (by acquiring a beneficial secondary alteration).

### Chromosome 1q aneuploidy suppresses p53 signaling by increasing MDM4 expression

We sought to uncover the biological mechanism underlying the addiction to chromosome 1q aneuploidy. RNA-seq analysis of A2780 cells revealed that elimination of the 1q-trisomy causes significant upregulation of tumor suppressor p53 target genes (Fig. 5A-B). Western blotting confirmed that 1q-disomic clones exhibit increased phosphorylation of p53 at serine-15 and increased expression of the canonical p53 target p21 (Fig. 5C)^55^. These results were not a by-product of CRISPR mutagenesis, as A2780 cells harboring a CRISPR-mediated integration of HSV-TK did not display evidence of p53 activation (Fig. 5B-C). Additionally, 1q-disomic cells exhibited a delay in the G1 phase of the cell cycle and increased levels of senescence-associated beta-galactosidase staining, both of which are associated with p53-mediated tumor suppression (Fig. S13)^56^. These results suggest that the chromosome 1q trisomy inhibits p53 signaling, and elimination of this trisomy antagonizes malignant growth at least in part by triggering p53 activation.

**Figure 5.**
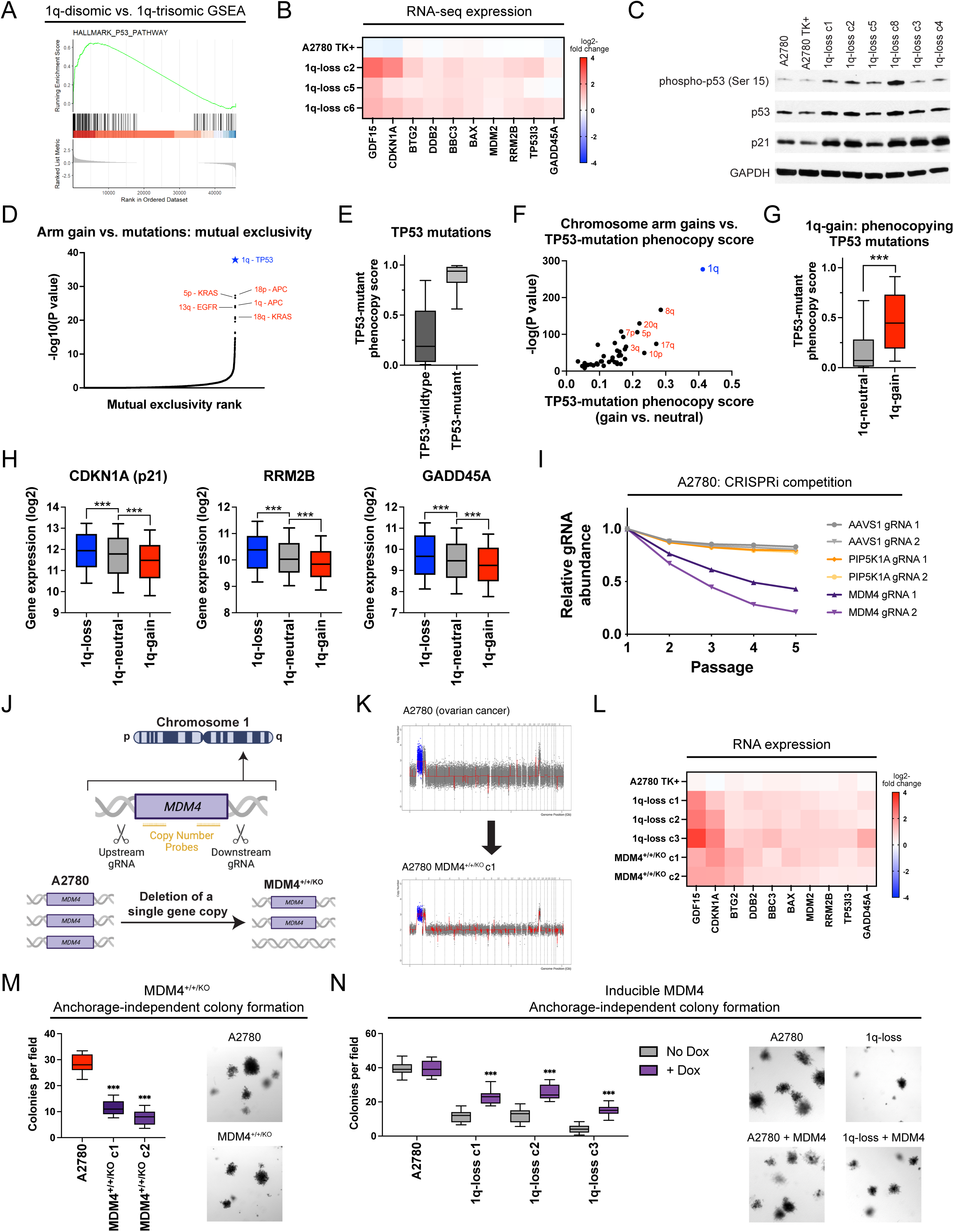
A single extra copy of MDM4 suppresses TP53 signaling and contributes to the 1q-trisomy addiction. (A) GSEA analysis of A2780 RNA-seq data reveals upregulation of the p53 pathway in the 1q-disomic clones, relative to the parental trisomy. (B) A heatmap displaying the upregulation of 10 p53 target genes in A2780 1q-disomic clones. The TK+ clone indicates a clone that harbors the CRISPR-mediated integration of the HSV-TK transgene but that was not treated to induce chromosome 1q-loss. (C) Western blot analysis demonstrating activation of p53 signaling in 1q-disomic clones. GAPDH was analyzed as a loading control. The TK+ clone indicates a clone that harbors the CRISPR-mediated integration of the HSV- TK transgene but that was not treated to induce chromosome 1q-loss. (D) A waterfall plot highlighting the most-significant instances of mutual exclusivity between chromosome arm gains and mutations in cancer-associated genes. The complete dataset for mutual exclusivity and co-occurrence is included in Table S1. (E) Boxplots displaying the TP53-mutation phenocopy signature^57^ in cancers from the TCGA, split based on whether the cancers harbor a non-synonymous mutation in TP53. (F) A scatterplot comparing the association between chromosome arm gains and the TP53-mutation phenocopy signature^57^ in TP53-wildtype cancers from TCGA. Cancers with chromosome 1q gains are highlighted in blue. (G) Boxplots displaying the TP53-mutation phenocopy signature^57^ in cancers from the TCGA, split based on whether tumors harbor a gain of chromosome 1q. Only TP53-wildtype cancers are included in this analysis. (H) Boxplots displaying the expression of three p53 target genes – CDKN1A (p21), RRM2B, and GADD45A – in cancers from TCGA split based on the copy number of chromosome 1q. Only TP53-wildtype cancers are included in this analysis. (I) A CRISPRi competition assay demonstrates that gRNAs targeting MDM4 drop out over time in A2780 cells. In contrast, gRNAs targeting AAVS1 and PIP5K1A, another gene encoded on chromosome 1q, exhibit minimal depletion. (J) A schematic displaying the strategy for using paired CRISPR gRNAs to delete a single copy of MDM4 in a cell line with a trisomy of chromosome 1q. (K) SMASH karyotype demonstrating maintenance of the chromosome 1q trisomy in an MDM4^+/+/KO^ clone. Chromosome 1q is highlighted in blue. (L) 1q-disomic clones and MDM4^+/+/KO^ clones in A2780 exhibit comparable upregulation of p53 transcriptional targets, as determined through TaqMan gene expression assays. (M) MDM4^+/+/KO^ clones exhibit decreased anchorage-independent growth relative to the MDM4^+/+/+^ parental cell line. (N) Induction of MDM4 cDNA in 1q-disomic clones in A2780 increases anchorage-independent growth. For the graphs in E, G, H, M, and N, the boxplots represent the 25th, 50th, and 75th percentiles of the indicated data, while the whiskers represent the 10th and 90th percentiles of the indicated data. For the soft agar experiments in M and N, the data are from n = 15 fields of view, and a representative trial is shown. ***p < 0.0005

To explore whether p53 inhibition is a common consequence of chromosome 1q gains, we examined our prior analysis of aneuploidy-mutation mutual exclusivity in cancer genomes (Table S1). Interestingly, out of 14,383 aneuploidy-gene mutation pairs, the single strongest instance of mutual exclusivity was between 1q gains and TP53 mutations (Fig. 5D). Put differently, while chromosome 1q gains and TP53 mutations are separately very common in cancer, individual tumors develop both 1q gains and TP53 mutations significantly less often than expected by chance (P < 10^-39^, Fisher’s exact test; Table S1). Next, we applied a recently-described classification algorithm capable of predicting cancers that lack p53 function based on their transcriptional profiles^57^. As expected, cancers from the TCGA with non-synonymous TP53 mutations scored significantly higher with this classifier than cancers with wild-type TP53 (Fig. 5E). Considering only tumors with wild-type TP53, we calculated the association between the p53 status classifier and every possible chromosome arm gain in the TCGA. Across all chromosomes, 1q gains exhibited the strongest correlation with the p53-loss signature (Fig. 5F-G). Among tumors with wild-type TP53, gains of chromosome 1q were associated with significantly lower expression of the p53 target genes CDKN1A (p21), GADD45A, and RRM2B (Fig. 5H)^55^. In total, these results indicate that gaining chromosome 1q phenocopies the effects of p53 mutations and suppresses p53 activity in human tumors.

We sought to discover the gene(s) on chromosome 1q responsible for inhibiting p53 signaling. We noted that MDM4, a known negative regulator of p53 activity, is located on 1q32^58^. MDM4 expression increased with chromosome 1q copy number and higher MDM4 expression correlated with the p53-loss transcriptional signature (Fig. S14A-B). To uncover whether MDM4 is directly responsible for the 1q-aneuploidy addiction observed in A2780, we first used CRISPR-interference (CRISPRi) to downregulate MDM4 expression without fully ablating it^59, 60^. In A2780 competition assays, we observed that downregulating MDM4 impaired cell fitness relative to A2780 cells in which AAVS1 or PIP5K1A, an unrelated gene on chromosome 1q, were downregulated (Fig. 5I)^61^. Next, we used a two-guide strategy to delete a single copy of MDM4 in an otherwise trisomic background (Fig. 5J-K and S14C-D). We found that the subsequent A2780 MDM4^+/+/KO^ clones downregulated MDM4 and upregulated TP53 target genes, comparable to the effects observed in cells lacking the entire 1q trisomy (Fig. 5L and S14E). We then tested the colony formation ability of MDM4^+/+/KO^ clones, and we discovered that losing a single copy of MDM4 significantly decreased anchorage-independent growth (Fig. 5M). Subsequently, we performed the converse experiment: we cloned MDM4 cDNA under the control of a doxycycline-inducible promoter and transduced the construct into both 1q-trisomic and 1q-disomic cells. Remarkably, we found that moderate (1.7-fold) overexpression of MDM4 was sufficient to cause a significant increase in anchorage-independent growth in the 1q-disomic cells, while this same treatment did not affect the 1q-trisomic cells (Fig. 5N and S14F).

Finally, to investigate the role of p53 as a mediator of 1q-aneuploidy addiction from an orthogonal approach, we used CRISPR to delete the TP53 gene in A2780 1q-disomic and 1q-trisomic cells (Fig. S15A). SMASH-Seq confirmed that these cells did not acquire any secondary chromosomal copy number changes during clonal derivation (Fig. S15B). We discovered that loss of TP53 rescued the G1 delay and significantly enhanced anchorage-independent growth in 1q-disomic cells (Fig. S15C-D). The magnitude of the increase in colony formation was significantly greater in the 1q-disomic cells compared to the 1q-trisomic cells (4-fold vs. 1.5-fold; P < .0001, t-test)(Fig. S15D). In total, these results indicate that chromosome 1q gains represent a mechanism by which p53-wildtype cancers can suppress p53 activity, and this suppression occurs due to the increased expression of MDM4. Nonetheless, we note that deleting TP53 and overexpressing MDM4 increased fitness in the 1q-disomic clones but did not fully restore fitness to wild-type levels. Thus, there are likely other dosage-sensitive fitness genes on chromosome 1q, and those genes may contribute to selection for 1q-gains in p53- mutant backgrounds.

### Chromosome 1q aneuploidy creates a collateral therapeutic vulnerability

The oncogene addiction hypothesis represents the conceptual foundation for the use of targeted therapies in cancer^51^. Cancers addicted to driver oncogenes like BCR-ABL or EGFR^L858R^ respond to inhibitors of those proteins that otherwise have minimal effect in untransformed tissue. Correspondingly, we sought to uncover whether aneuploidy addictions could also represent a therapeutic vulnerability for certain cancers. Toward that goal, we noted that chromosome 1q harbors the UCK2 gene, which encodes a kinase involved in the pyrimidine salvage pathway^62^. UCK2-dependent phosphorylation has previously been reported to function as the rate-limiting step in the metabolism of certain nucleotide analogs, including RX-3117 and 3-deazauridine (Fig. 6A)^63–65^. Phosphorylated RX-3117 and 3-deazauridine can poison cellular nucleotide pools and block DNA and RNA synthesis^66^. We found that UCK2 is over-expressed in human cancers that contain extra copies of chromosome 1q, and elimination of the chromosome 1q trisomy consistently decreased UCK2 protein expression in our engineered cell lines (Fig. 6B-C). We therefore investigated whether gaining chromosome 1q could create a collateral sensitivity to UCK2-dependent nucleotide analogs.

**Figure 6.**
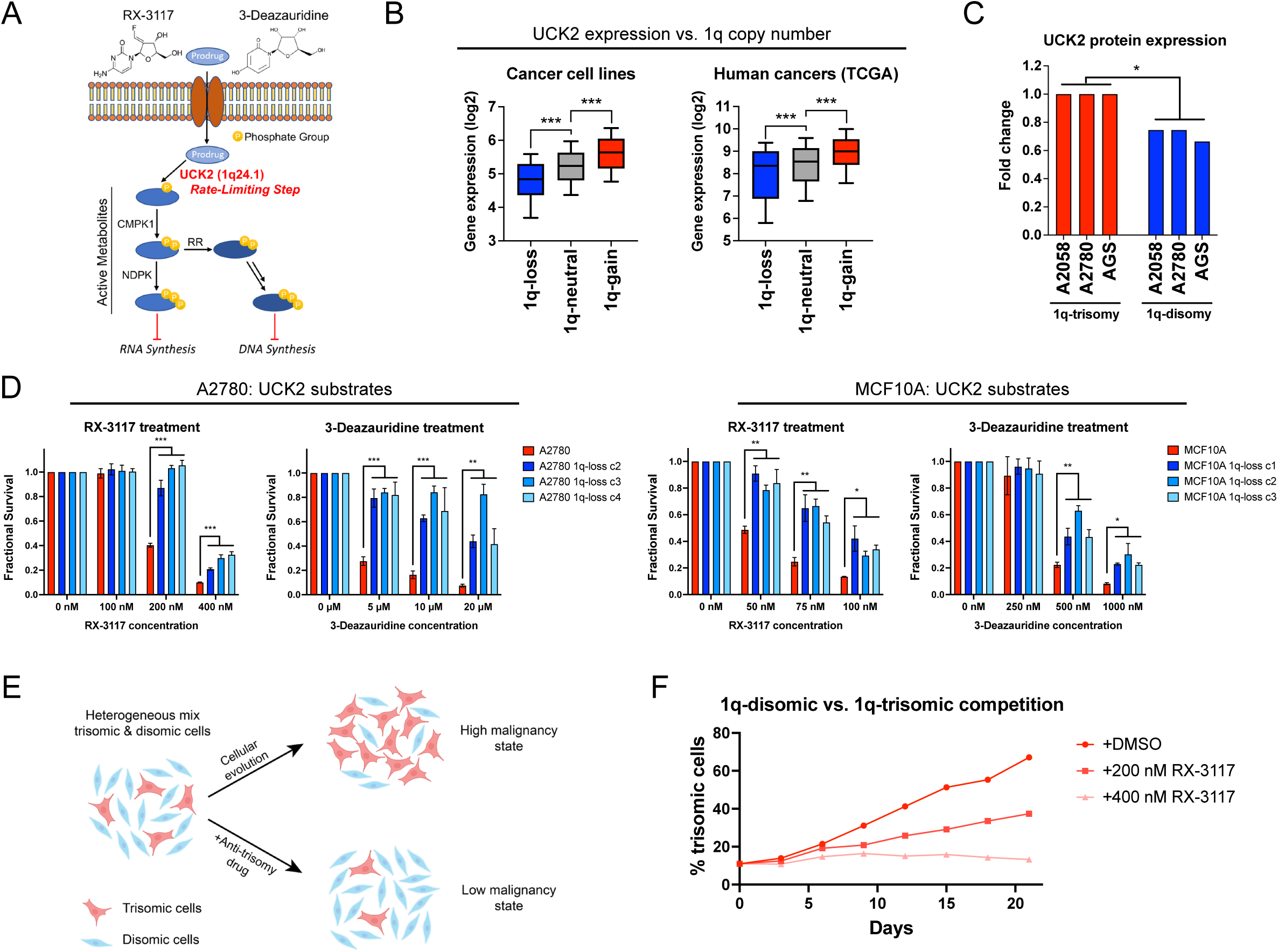
Gaining chromosome 1q increases sensitivity to UCK2 substrates. (A) A schematic of the metabolism of two pyrimidine analogs, RX-3117 and 3-deazauridine. UCK2, a kinase encoded on chromosome 1q, phosphorylates these compounds to produce cytotoxic derivatives that can poison DNA and RNA synthesis. (B) Boxplots displaying the expression of UCK2 in cancer cell lines^126^ (left) and human cancers^128^ (right), divided based on the copy number of chromosome 1q. The boxplots represent the 25th, 50th, and 75th percentiles of the indicated data, while the whiskers represent the 10th and 90th percentiles of the indicated data. (C) Expression of UCK2 protein in cancer cell lines with 1q trisomies or following aneuploidy-elimination. (D) Cellular sensitivity of A2780 and MCF10A treated with different concentrations of RX-3117 or 3-deazauridine. (E) A schematic displaying cellular competition between trisomic and disomic cells. Under normal conditions, certain trisomies enhance cellular fitness, allowing these cells to overtake the population and enhance malignant growth (top). However, treatment with an “anti-trisomy” compound could selectively impair the growth of the aneuploid cells, keeping the population in a low-malignant state (bottom). (F) A cellular competition between fluorescently-labeled A2780 1q-trisomic and unlabeled 1q-disomic cells. These cells were mixed at a ratio of 10% to 90% and then cultured in either DMSO or RX-3117. While the trisomic cells quickly dominate the population in drug-free media, treatment with RX-3117 prevents the outgrowth of the 1q-trisomy subpopulation. *p < 0.05, ** p < 0.005, *** p < 0.0005

First, as the mechanism of many cancer drugs is poorly-understood^67, 68^, we sought to verify that the cytotoxicity of RX-3117 and 3-deazauridine requires UCK2 expression. We used CRISPR to delete UCK2 in the haploid HAP1 cell line, and we confirmed that UCK2-knockout cells are highly resistant to both compounds (Fig. S16A-C). Next, we tested the effects of RX-3117 and 3-deazauridine in our engineered 1q-trisomic and 1q-disomic cell lines. We found that A2780 and MCF10A cells harboring a trisomy of chromosome 1q were significantly more sensitive to both compounds compared to isogenic cells containing two copies of chromosome 1q (Fig. 6D). This effect was specific for UCK2 substrates, as the 1q-trisomic cells did not exhibit greater sensitivity to UCK2- independent nucleotide poisons and other cancer drugs (Fig. S16D). Furthermore, ectopic over-expression of UCK2 cDNA in 1q-disomic clones was sufficient to re-sensitize these cells to RX-3117 (Fig. S16E). Finally, RX- 3117 and 3-deazauridine have previously been screened across the NCI-60 cell line panel, and we found that higher UCK2 expression levels correlate with greater sensitivity to both compounds (Fig. S16F)^69^. In total, these results indicate that 1q gains induce a collateral sensitivity to specific nucleotide analogs by increasing the expression of the UCK2 kinase.

We hypothesized that we could use the greater sensitivity of 1q-trisomic cells to UCK2 substrates to re-direct cellular evolution away from aneuploidy and towards a disomic state with lower malignant potential (Fig. 6E). We mixed fluorescently-labeled 1q-trisomic and 1q-disomic A2780 cells at a ratio of 10:90, and then co-cultured the two cell populations. After three weeks of growth in drug-free media, the trisomic cells had expanded to make up 67% of the culture, consistent with our observation that the extra copy of chromosome 1q enhances cellular fitness (Fig. 6F). In contrast, when the same cell populations were grown in the presence of 400 nM RX-3117, there was no significant increase in the trisomic cell population over time. We conclude that trisomy-selective compounds can be used to manipulate cellular evolution, potentially providing a new strategy to prevent the outgrowth of malignant aneuploid cells in a pre-malignant setting.

## DISCUSSION

Here, we describe ReDACT, a set of tools that can be used to eliminate aneuploid chromosomes from human cell lines. Using ReDACT, we engineered genetically-matched cell lines that have or lack common cancer aneuploidies, and we demonstrate that losing the aneuploidy of chromosome 1q blocks malignant growth in cell lines harboring this alteration. We posit that these phenotypes are due specifically to the loss of the aneuploid chromosome and are not a by-product of CRISPR selection or the elimination of point mutations encoded on the targeted chromosome (discussed in more detail in Supplemental Text 1 and 2, respectively). Due to the similarity between our observations and the previously-described cancer gene addiction phenomenon, we suggest that, in certain circumstances, cancers may also be “addicted” to the aneuploidy found in their genomes.

Historically, cancer aneuploidy has been resistant to close analysis^5^. While individual cancer driver genes have been studied for several decades using the standard tools of molecular genetics, manipulating chromosome dosage has been technically challenging. Initial insight into the role of aneuploidy in tumorigenesis was gained through the development of genetically-engineered mouse models that harbor mutations in mitotic checkpoint genes^70–72^. These chromosomal instability-promoting mutations were found to either enhance or suppress tumorigenesis, depending on the oncogenic stimulus and the degree of instability^73–75^. However, it is not clear whether these results can be attributed specifically to aneuploidy itself, as chromosome missegregation can cause certain phenotypes that are independent of the resulting dosage imbalance^76–78^ and many mitotic checkpoint genes moonlight in other cellular processes^79–81^. Alternately, microcell-mediated chromosome transfer has been used to introduce extra chromosomes into diploid cell lines^82–84^. Cells manipulated to carry trisomic chromosomes were found to exhibit reduced malignant potential compared to their diploid parental cell lines^85^. These tumor-suppressive effects of aneuploidy have been attributed to the global imbalance in protein stoichiometry caused by the expression of hundreds of extra genes^86–89^.

In this work, we eliminated endogenous aneuploidies from established cancer cell lines, and we revealed that loss of these trisomic chromosomes compromised cancer-like growth. We posit that during tumor evolution, certain aneuploidies can provide context-specific benefits that enhance tumorigenesis. For instance, we showed that chromosome 1q gains are an early event during breast cancer development, and we demonstrated that MDM4 is a dosage-sensitive gene on 1q that suppresses p53 signaling and enhances cancer growth. In other cancer types or in cells that already harbor TP53 mutations, the beneficial effects of gaining MDM4 may be outweighed by the detrimental effects of overexpressing hundreds of additional 1q genes.

We note that MDM4 and many other genes have been demonstrated to have tumor-promoting properties when highly overexpressed^90^. For instance, MDM4 is focally amplified in ∼1% of cancers in the TCGA, and strong overexpression of MDM4 via retrovirus immortalizes primary cells and renders them sensitive to RAS-mediated transformation^91^. Our results demonstrate that even a single extra copy of MDM4 can be oncogenic, and we speculate that there are many genes, both known and unknown, that can promote tumorigenesis when their copy number is increased from two to three. The overlap between these single-copy dosage-sensitive genes and genes found to have tumor-promoting ability when highly overexpressed is at present unknown.

Finally, our results raise the exciting possibility that “aneuploidy addictions” may represent a new therapeutic vulnerability in cancer. Previous attempts to target aneuploidy have focused on phenotypes that are shared across highly-aneuploid cells, such as alterations in spindle geometry^92, 93^. Here, we sought to develop an approach to take advantage of the genes that are encoded on an aneuploid chromosome, thereby allowing chromosome-specific targeting. In particular, we hypothesized that the over-expression of specific genes – for instance, drug-importer pumps or enzymes required for a pro-drug’s activation – could sensitize cancers to compounds that are otherwise better-tolerated in euploid tissue. We demonstrated that gaining chromosome 1q creates a collateral vulnerability to the nucleotide analogs RX-3117 and 3-deazauridine due to the overexpression of the kinase UCK2. Notably, RX-3117 has been tested in phase 2A clinical trials, but without the use of any genomic biomarkers for patient selection^94^. We speculate that this drug may be particularly effective if given to patients with tumors that harbor gains of chromosome 1q. More broadly, compounds whose anti-cancer function is enhanced by genes encoded on aneuploid chromosomes could be used to direct cellular evolution away from certain aneuploidies and toward the lower-malignancy diploid state.

## ACKNOWLEDGMENTS

We are grateful to Dr. Tobias Cantz (Hannover Medical School) for providing the TK plasmids used in this work. We thank Peter Andrews (CSHL) for assistance with SMASH-Seq. We thank Yale Flow Cytometry, especially Chao Wang and Lesley Devine, for their assistance with single-cell sorting. Yale Flow Cytometry is supported in part by an NCI Cancer Center Support Grant # NIH P30 CA016359. We thank Al Mennone and the Yale Center for Advanced Light Microscopy Facility for their assistance with soft agar imaging. We thank the Yale Center for Genome Analysis for performing SMASH-Seq. We thank the Yale Center for Research Computing, specifically Robert Bjornson, for guidance and assistance in computation run on the Farnam and Ruddle clusters. We thank the Yale Animal Resources Center Staff for assistance with mouse experiments. This work was performed with assistance from the CSHL Flow Cytometry, Microscopy, Animal, and Sequencing Technologies & Analysis Shared Resources, which are supported in part by the Cancer Center Support Grant 5P30CA045508.

Copy number timing analysis conducted in the Sun Lab uses the computing resources of the Minnesota Supercomputing Institute. The timing analysis was prepared using limited access datasets obtained from the Cancer Genome Project from the Wellcome Sanger Institute and does not necessarily reflect the opinions of the provider institution. Part of the BRCA sequencing data was originally generated by research led by Dr. Masahito Kawazu and is available at the website of the National Bioscience Database Center (NBDC; http://biosciencedbc.jp/en/) of the Japan Science and Technology Agency (JST). We also thank the International Cancer Genome Consortium (ICGC) for providing access to the MEL dataset.

Research in the Sheltzer Lab is supported by NIH grant R01CA237652, Department of Defense grant W81XWH- 20-1-068, an American Cancer Society Research Scholar Grant, a Breast Cancer Alliance Young Investigator Award, a Damon Runyon-Rachleff Innovation Award, a sponsored research agreement from Ono Pharmaceuticals, and a sponsored research agreement from Meliora Therapeutics. Research in the Liu Lab is supported by NIH grant R01GM137031.

## MATERIALS & METHODS

### BASIC CELL CULTURE TECHNIQUES

#### Cell lines and culture conditions

The identities of all cell lines used in this study were confirmed using STR (short tandem repeat) profiling (University of Arizona Genetics Core). A2780 was grown in RPMI 1640 medium (Gibco, cat. no. 11875119) supplemented with 10% fetal bovine serum (FBS) (Sigma-Aldrich, cat. no. F4135), 2 mM glutamine (Lonza, cat. no. 17-605F), and 100 U/mL penicillin-streptomycin (Life Technologies, cat. no. 15140122). A2058, HCT116, HEK-293T, and PLAT-A cells were grown in DMEM (Gibco, cat. no. 11995073) supplemented with 10% FBS, 2 mM glutamine, and 100 U/mL penicillin-streptomycin. AGS was grown in F-12K (ATCC; cat. no. 30-2004) supplemented with 10% FBS, 2 mM glutamine, and 100 U/mL penicillin-streptomycin. HAP1 cells were grown in IMDM (Gibco, cat. no. 12440053) supplemented with 10% FBS, 2 mM glutamine, and 100 U/mL penicillin-streptomycin. MCF-10A was grown in DMEM/F-12 (Gibco, cat. no. 11320082), supplemented with 5% horse serum (Gibco, cat. no. 16050122), 20 ng/mL EGF (PeproTech, cat. no. AF-100-15), 0.5 ng/mL Hydrocortisone (Sigma-Aldrich, cat. no. H0888), 100 ng/mL Cholera Toxin (Sigma-Aldrich, cat. no. C8052), 10 µg/mL Insulin (Sigma-Aldrich, cat. no. I1882), 5 mM Transferrin (Sigma-Aldrich, cat. no. T8158), and 100 U/mL penicillin-streptomycin. All cells were cultured in a humidified environment at 37°C and 5% CO2. Sources of the cell lines used in this manuscript are listed in Table S4.

#### Production of lentivirus and retrovirus

HEK293T (lentivirus) or PLAT-A (retrovirus) cells were transfected using the calcium-phosphate method^95^. Virus-containing supernatant was harvested 48 to 72 hours post-transfection, filtered through a 0.45 μm syringe, and then frozen at -80°C for later use or applied directly to cells with 4-10 μg/mL polybrene (Santa Cruz Biotechnology, Inc., cat no. SC-134220). The culture media on target plates was changed 24 hours post-transduction.

### PLASMID CLONING METHODS

#### CRISPR plasmid cloning

Guide RNAs for CRISPR experiments were designed with Benchling (www.benchling.com). Guides were cloned into the Lenti-Cas9-gRNA-GFP vector (Addgene #124770) or LRCherry2.1 (Addgene #108099) using a BsmBI digestion as previously described^96^. Plasmids were transformed in Stbl3 E. coli (Thermo Fisher Scientific, cat. no. C737303) and sequence-verified to confirm the presence of the correct gRNA. CRISPR gRNA sequences are listed in Table S5A.

#### CRISPRi plasmid cloning

Guide RNAs for CRISPRi experiments were chosen from refs^60, 97^, and two guides per gene were cloned into the LRG2.1 mCherry vector (Addgene #108099) as described above. Plasmids were transformed in Stbl3 *E. coli* and sequenced to confirm the presence of the correct gRNA sequence. CRISPRi gRNA sequences are listed in Table S5B.

#### Cloning of doxycycline-inducible MDM4

The MDM4 coding sequence (NM_002393.5) was cloned into Lenti-X™ Tet-One™ Inducible Expression System (Takara Bio, cat. no. 631847). The resulting plasmid was then subject to whole-plasmid sequencing for verification. A2780 cells were transduced with pLVX-TetOne-MDM4-Puro and selected for with 1.5 μg/mL puromycin (InvivoGen, cat. no. ant-pr-1). Tight inducibility was confirmed through doxycycline (Sigma-Aldrich, cat. no. D3072) titration and quantitative real-time PCR. The final plasmid was deposited on Addgene (#195140).

### KARYOTYPING AND DNA COPY NUMBER ANALYSIS

#### SMASH-seq

Libraries for genomic copy number analysis were prepared as described in ref^43^. In brief, genomic DNA was enzymatically fragmented to a mean size of ∼40bp and ligated to generate long chimeric DNA molecules (∼300- 700bp) for sequencing. Fragment size selection and purification were done with Agencourt AMPure XP beads (Beckman Coulter, cat. no. A63881). Illumina-compatible NEBNext Multiplex Dual Index primer pairs and adapters (New England Biolabs, cat. no. E6440S) were added to each sample, and the products were pooled for next-generation sequencing (NGS). Libraries were sequenced using an Illumina MiSeq or NovoSeq sequencer. The generated reads were demultiplexed then mapped using a custom Nextflow^98^ wrapper running *SMASH* built from MUMdex^99^ (commit: 25e1f2f). Plots were generated via a custom script utilizing *tidyverse*^100^ (v1.3.1) components run in the R programming environment (R Core Team 2022; v4.2.0).

#### G-banding karyotyping

A2780 samples were sent to Cell Line Genetics Inc. (www.clgenetics.com), and AGS and A2058 samples were sent to Karyologic Inc. (www.karyologic.com) for G-banding karyotyping. A minimum of 10 metaphase spreads per sample were counted and analyzed to prepare representative karyotype spreads.

#### TaqMan copy number analysis

Genomic DNA was extracted and isolated using the Qiagen DNeasy Blood & Tissue kit (Qiagen, cat. no. 69506). Reactions were prepared in quadruplicate, and target probes were duplexed with RNaseP (Applied Biosystems, cat. no. 4316831) or TERT (Applied Biosystems, cat. no. 4403316) as the reference assay. Quantitative PCR was performed using TaqPath ProAmp Master Mix (Applied Biosystems, cat. no. A30867) and quantified using the QuantStudio 6 Flex Real-Time PCR system (Applied Biosystems). Copy number analysis was performed as described in ref^101^ and copy number calls were normalized to the near-diploid colorectal cancer cell line DLD1^102^. To screen for chromosome arm loss, a minimum of three probes spanning the chromosome of interest were used, and the copy number calls for the individual probes were averaged to determine chromosome arm copy number. To screen for single copy gene deletions, two probes for the gene of interest were used, and the copy number calls for the individual probes were averaged to determine gene copy number. TaqMan copy number probes are listed in Table S6.

### GENE EXPRESSION ANALYSIS

#### RNAseq

Total cellular RNA was extracted using the Qiagen RNeasy Mini Kit (Qiagen, cat. no. 74106) and subjected to on-column DNase digestion (Qiagen, cat. no. 79254). Purified RNA samples were submitted to Novogene for RNAseq and quantification. In brief, mRNA was purified from total cellular RNA using poly-T oligo-attached magnetic beads. After fragmentation, first strand cDNA was synthesized using random hexamer primers, followed by second strand cDNA synthesis. Subsequently, libraries were prepared with end repair, A-tailing, adapter ligation, size selection, amplification, and purification. Libraries were sequenced on an Illumina platform and paired-end reads were generated. Raw data (raw reads) of fastq format were firstly processed through in-house Novogene Perl scripts. Reference genome and gene model annotation files were downloaded from genome website directly. An index of the reference genome was built using Hisat2 v2.0.5 and paired-end clean reads were aligned to the reference genome using Hisat2 v2.0.5. featureCounts v1.5.0-p3 was used to count the reads numbers mapped to each gene. The FPKM of each gene was calculated based on the length of the gene and reads count mapped to this gene. Gene Set Enrichment analysis was performed with the local version of the GSEA analysis tool http://www.broadinstitute.org/gsea/index.jsp and the predefined Hallmark gene sets.

#### Mass spectrometry

Proteomic analysis was conducted as previously described^103^. In short, cell pellets were thawed and a VialTweeter device (Hielscher-Ultrasound Technology) was used to sonicate the samples (4 °C; 1 min; two cycles). The samples were centrifuged at 20,000 g for 1 hour to remove insoluble material. Protein concentration was measured using the Bio-Rad protein assay dye (Bio-Rad, cat. no. 5000006). 800 µg of protein per sample were diluted (final concentration = 2 µg/µL) using a 10 M urea/100 mM ammonium bicarbonate buffer, reduced by 10 mM DTT (1 hour; 56 °C), and alkylated by 20 mM IAA (1 hour; RT). The proteins were subjected to a precipitation-based digestion^104^. Briefly, five volumes of precooled precipitation solution (50% acetone, 50% ethanol, and 0.1% acetic acid) were added to the samples. After overnight incubation at -20 °C, the samples were centrifuged (20,000 x g; 4 °C; 40 min). The precipitate was washed with precooled 100% acetone, centrifuged (20,000 x g; 4 °C; 40 min), and the remaining acetone was evaporated in a SpeedVac. For protein digestion, 300 µL of 100 mM NH4HCO3 with sequencing grade porcine trypsin (Promega) at a trypsin-to-protein ratio of 1: 20 were added and incubated overnight at 37 °C. The resulting peptide samples were acidified with formic acid and desalted using a C18 column (MarocoSpin Columns, NEST Group INC.) according to the manufacturer’s instructions.

1 µg of the peptide mixture was used for the LC-MS analysis as described previously^103, 105^. The LC separation was performed using an EASY-nLC 1200 system (Thermo Scientific) using a self-packed PicoFrit column (New Objective, Woburn, MA, USA; 75 µm × 50 cm length) with a ReproSil-Pur 120A C18-Q 1.9 µm resin (Dr. Maisch GmbH, Ammerbuch, Germany). A 120-min gradient length was used to elute peptides from the LC; with buffer B (80% acetonitrile containing 0.1% formic acid) from 5% to 37% and corresponding buffer A (0.1% formic acid in H2O). The flow rate was 300 nL/ min, and the temperature was controlled at 60 °C using a column oven (PRSO-V1, Sonation GmbH, Biberach, Germany). The Orbitrap Fusion Lumos Tribrid mass spectrometer (Thermo Scientific) coupled with a NanoFlex ion source (spray voltage of 2000 V, 275 °C) was used for the MS analysis. The method for DIA-MS consisted of a MS1 survey scan and 33 MS2 scans of variable windows^106, 107^. The MS1 scan parameters were set as follows: scan range 350–1650 m/z, resolution 120,000 at m/z 200, the AGC target 2.0E6, and the maximum injection time 100 ms. The normalized HCD collision energy was 28%. The MS2 scan parameters were the following: resolution 30,000 at m/z 200, the AGC target 1.5E6, and the maximum injection time 50 ms. The default peptide charge state was set to 2. Both of MS1 and MS2 spectra were recorded in a profile mode.

The DIA-MS data analysis was performed using Spectronaut v15^106, 108^ using the library-free DirectDIA workflow^106, 109^ and the Swiss-Prot protein database (September 2020, 20,375 entries). The analysis was performed using default Spectronaut settings. Methionine oxidation and N-terminal acetylation were set as variable modifications, where carbamidomethylation at cysteine was set as a fixed modification. Both peptide- and protein-FDR were controlled at 1%, and the resulting data matrix was filtered by “Qvalue”. The DIA quantification was performed using the MS2 level peak areas. Protein intensities were exported, log2- transformed and normalized using LOESS normalization^110^ prior to the subsequent analysis.

#### Quantitative real-time PCR

Total cellular RNA was extracted and isolated using the Qiagen RNeasy Mini Kit (Qiagen, cat. no. 74106). cDNA synthesis was performed using SuperScript IV VILO Master Mix (Invitrogen, cat. no. 11756500). Quantitative PCR was performed for the target genes using TaqMan Fast Advanced Master Mix (Applied Biosystems, cat. no. 4444963) and quantified using the QuantStudio 6 Flex Real-Time PCR system (Applied Biosystems). TaqMan gene expression assays are listed in Table S7 and qPCR primers are listed in Table S8.

#### Western Blotting

One day prior to lysate harvest, 500,000 cells were seeded in a six-well plate. Whole cell lysates were harvested and resuspended in RIPA buffer [25 mM Tris, pH 7.4, 150 mM NaCl, 1% Triton X 100, 0.5% sodium deoxycholate, 0.1% sodium dodecyl sulfate, protease inhibitor cocktail (Sigma, cat. no. 4693159001), and phosphatase inhibitor cocktail (Sigma, cat. no. 4906845001)]. Protein concentration was quantified using the RC DC Protein Assay (Bio-Rad; cat. no. 500–0119). Equal amounts of lysate were denatured and loaded onto a 10% SDS-PAGE gel. The Trans-Blot Turbo Transfer System (Bio-Rad) and polyvinylidene difluoride membranes were used for protein transfer. Antibody blocking was done with 5% milk in TBST (19 mM Tris base, NaCl 137 mM, KCl 2.7 mM and 0.1% Tween-20) for 1 hour at room temperature. The following antibodies and dilutions were used: p21 (Abcam; cat. no. ab109520) at a dilution of 1:1000 in 5% milk, p53 (Abcam; cat. no. ab1101) at a dilution of 1:1000 in 5% milk, p53 (Abcam; cat. no. ab32389) at a dilution of 1:1000 in 5% milk, Phospho-p53 (Ser15) (Cell Signaling, cat. no. 9284S) at a dilution of 1:1000 in 5% milk, Cleaved-PARP (Asp214) (Cell Signaling; cat. no. 5625S) at a dilution of 1:1000 in 5% milk, and UCK2 (proteinTech; cat. no. 10511-1-AP) at a dilution of 1:1000 in 5% milk. Blots were incubated with the primary antibody overnight at 4°C. Anti-GAPDH (Santa-Cruz; cat. no. sc-365062) at a dilution of 1:20,000 in 5% milk, or Anti-alpha tubulin (Sigma-Aldrich; cat. no. T6199) at a dilution of 1:20,000 in 5% milk was used as a loading control. Membranes were washed at room temperature three times (20 mins each) before they were incubated in secondary antibodies for an hour at room temperature. HRP goat anti-mouse (Bio-Rad; cat. no. 1706516) at 1:20,000 was used for tubulin, p53 (ab1101), MDM4, and GAPDH blots while HRP goat anti-rabbit (Abcam; cat. no. ab6721) at 1:20,000 was used for all other primary antibodies. Membranes were washed three times again (20 min each) and developed using ProtoGlow ECl (National Diagnostics; cat. no. CL-300) and autoradiographic film (Lab Scientific; XAR ALF 2025).

### CHROMOSOME ENGINEERING

#### Chromosome elimination: ReDACT-NS

##### Generation of selection cassette

A centromere-proximal CRISPR gRNA was designed and cloned into the Lenti-Cas9-gRNA-GFP vector (Addgene #124770) for integration of the selection cassette. Homology arms for cassette integration were designated as the 180bp immediately upstream and downstream of the guide RNA targeting site. Adapters for PCR were added to the 3’ end of the 180bp homology arms (Forward: gacattgattattgactagt; Reverse: ccatagagcccaccgcatcc), and the resulting 200bp ultramers were obtained from IDT. PCR for selection cassette production was performed with SeqAmp DNA Polymerase (Takara Bio, cat. no. 63850) using the ultramers as primers and AAT-PB-CD2APtk (Addgene #86004) or AAT-PB-PG2APtk^111^ (Addgene #195124) as the template. The PCR products were purified and concentrated with the QIAquick PCR Purification kit (Qiagen, cat. no. 28106). Homology arm and ultramer sequences are listed in Table S8.

##### Knocking-in cassettes with CRISPR-mediated HDR

Cells were transfected with the PCR-purified selection cassette and integration gRNA CRISPR plasmid, using Lipofectamine 3000 (Invitrogen, cat. no. L3000015) for A2780, AGS & HCT116 cells, or FuGENE HD (Promega, cat. no. E2311) for A2058 cells. Integration of the cassette was selected with puromycin (InvivoGen, cat. no. ant-pr-1). Following selection, dsRed positive cells (when using Addgene #86004) or GFP positive cells (when using Addgene #195124) were single cell sorted onto 96-well plates, and clonal cell lines were established. Successful cassette integration was confirmed through PCR and sequencing. PCR check primer sequences are listed in Table S8. Integrant clones were subject to karyotypic validation through SMASH karyotyping prior to inducing chromosome arm loss.

##### Inducing chromosome arm loss

A centromere targeting CRISPR gRNA was designed and cloned into the Lenti-Cas9-gRNA-GFP vector (Addgene #124770) for targeted chromosome elimination. Integrant clones were transfected with the centromere targeting gRNA CRISPR plasmid using Lipofectamine 3000 or FuGENE HD. Chromosome arm loss was selected for with 10 µM ganciclovir (Sigma Aldrich, cat. no. G2536). Following ganciclovir selection, dsRed or GFP negative cells were single cell sorted onto 96-well plates, and clonal cell lines were established. Clones were screened for targeted chromosome arm loss with TaqMan copy number assays as described below and chromosome elimination was confirmed through SMASH karyotyping. The aneuploidy-loss cell lines generated using this method are listed in Table S3.

#### Chromosome elimination: ReDACT-TR

##### Generation of the artificial telomere construct

The ReDACT-TR technique was motivated by the use of a telomere seed sequence to generate monosomic cells as described in ref^1^. To enhance the efficacy of this approach, we created new plasmids linking a telomere seed sequence with a puromycin selection marker, which allowed us to enrich for stably-transfected aneuploidy-loss cells using drug selection. These EF1a-Puro-Telo vectors (Addgene #195138 and #195139) were generated by introducing a puromycin selection marker to a telomere seed sequence gifted by Alison Taylor. This vector was digested with FastDigest KpnI (Thermo Fisher Scientific, cat. no. FD0524) and FastDigest BstZ17I (Thermo Fisher Scientific, cat. no. FD0704), and gel purified to obtain the artificial telomere construct.

##### Inducing chromosome arm-loss with CRISPR-mediated NHEJ

Cells were transfected with the purified artificial telomere construct and centromere targeting gRNA CRISPR plasmids using Lipofectamine 3000 for AGS cells, or Fugene HD for MCF10A cells. Telomere replacement was selected for with puromycin, and puromycin resistant cells were single cell sorted onto 96-well plates. Clonal cell lines were established and screened for targeted chromosome arm loss with TaqMan copy number assays. Chromosome elimination was confirmed through SMASH karyotyping. The aneuploidy-loss cell lines generated using this method are listed in Table S3.

#### Chromosome elimination: ReDACT-CO

Cells were transfected with centromere targeting gRNA CRISPR plasmids using Lipofectamine 3000 or FuGENE HD. GFP+ cells were single cell sorted onto 96-well plates. Clonal cell lines were established and screened for targeted chromosome arm loss with TaqMan copy number assays. Chromosome elimination was confirmed through SMASH karyotyping. Chr1q_Centromere_Targeting_gRNA (Addgene #195125) and Chr1q_Cassette- Integration_gRNA (Addgene #195126) were used together to target chromosome 1q, Chr7_Centromere- Targeting_gRNA (Addgene #195129) was used to target chromosome 7p, and Chr8q_Centromere- Targeting_gRNA (Addgene #195128) was used to target chromosome 8q (Table S5A). The aneuploidy-loss cell lines generated using this method are listed in Table S3.

#### Choice of ReDACT techniques for each cell line

No underlying reasons motivated the selection of a specific ReDACT technique for each cell line. Our first approach, ReDACT-NS, was driven by the historical use of thymidine kinase for genomic modification in embryonic stem cells^112, 113^. However, after this project was initiated, several reports were published demonstrating accidental CRISPR-mediated chromosome loss without drug selection^41, 42^, which prompted us to experiment with the ReDACT-TR and ReDACT-CO techniques. Consequently, for most cell lines described in this manuscript, we initiated multiple ReDACT approaches at the same time, and then focused on characterizing the clones in which we were able to verify aneuploidy-loss first. Over time, we discovered that each technique had certain benefits and drawbacks. For instance, with ReDACT-NS, once a clone was isolated containing on-target integration of the HSV-TK cassette, it was relatively straightforward to produce a large number of aneuploidy-loss clones. In contrast, ReDACT-CO required only one round of single-cell cloning and was much faster than ReDACT-NS, but the overall efficiency of ReDACT-CO tended to be lower than ReDACT-NS. Importantly, we obtained consistent results on the effects of aneuploidy-loss regardless of the ReDACT technique that was ultimately used, underscoring our conclusion that aneuploidy-loss itself is the root cause of the observed phenotypes.

### PHENOTYPIC CHARACTERIZATION OF ANEUPLOIDY-LOSS CELLS

#### Proliferation assays

Cells were seeded in 6-well plates at 100,000 cells per well. After 72 hours, cells were harvested, counted, and 100,000 cells were re-plated in fresh media on 6-well plates. Cells were passaged ten times, and population doublings were calculated at each passage.

#### Soft agar assays

To assay colony formation in soft agar, a solution of 1.0% Difco Agar Noble (VWR Scientific, USA, cat. no. 90000–772) in sterile water was prepared. The 1% agar solution was mixed 1:1 with the base cell culture media supplemented with 20% FBS, 4 mM glutamine, and 200 U/mL penicillin/streptomycin, or for MCF10A, supplemented with 10% horse serum, 40 ng/mL EGF, 1 ng/mL Hydrocortisone, 200 ng/mL Cholera Toxin, 20 µg/mL Insulin, 10 mM Transferrin, and 200U/mL penicillin streptomycin. 1 mL of this mixture was plated on each well of a 6-well plate and allowed to solidify at room temperature to form a base layer of 0.5% agar. Cells were then harvested and counted. For A2780 and A2058, 20,000 cells were seeded in 0.35% agar (1:1 mixture of 0.7% agar in sterile water and 2x supplemented growth medium). For AGS and MCF10A, 10,000 cells were seeded in 0.35% agar. For HCT116, 35,000 cells were seeded in 0.30% agar. Plates were left at room temperature to solidify and then placed in a humidified incubator at 37°C and 5% CO2. 1 mL of normal growth media was added the next day, and every three days after. After 10-14 days, cells were fixed with 100% methanol, and stained with 0.01% crystal violet dissolved in 25% methanol. Colony formation was quantified by capturing z-stacks of several fields of view per well from at least three wells on a LSM 710 confocal microscope (Zeiss) or a Cytation5 imaging reader (BioTek Instrument, inc.), under either 4x, 5x, or 6x magnification. The average number of colonies was calculated by counting total number of colonies per field of view from multiple fields of view from at least three wells.

#### Xenograft assays

To assay tumor formation, cells were harvested and resuspended in cold PBS. For A2780 xenograft experiments, 3 million cells were injected in each flank of NU/J mice (Jackson Laboratory, cat. no. 002019). For A2058 xenograft experiments, 2 million cells were injected in each flank in NU/J mice. For HCT116 xenograft experiments, 4 million cells were injected in J:NU mice (Jackson Laboratory, cat. no. 007850). For MCF10A cells expressing HRAS^G12V^, 10 million cells were injected in each flank in NU/J mice. For AGS xenograft experiments, the following conditions were tried: 5 million cells resuspended in PBS in each flank of NU/J mice (Jackson Laboratory, cat. no. 002019); 4 million cells resuspended in PBS in each flank of NGS mice (Jackson Laboratory, cat. no. 005557); and 15 million cells resuspended in a 1:1 PBS:Matrigel mixture in each flank of J:NU mice (Jackson Laboratory, cat. no. 007850). Cells were subcutaneously injected using a 1 mL 25G x 5/8 syringe (BD, cat. no. 309626). Mice were visually monitored for tumor formation routinely following injection. Once a tumor was visible, it was measured every three days by calipers. Tumor volume was calculated using the formula V = ½ (longer axis)(shorter axis). All mouse protocols were approved by the CSHL and Yale Institutional Animal Care and Use Committees.

#### Derivation and characterization of cell lines post-xenograft

To derive post-xenograft cell lines, mice were euthanized at a humane end point, and tumors cut out using a sterilized pair of scissors. The tumors were placed into a chilled conical tube containing 2-3 mL of TrypLE Express Enzyme (Gibco; cat. no. 1260413) and transported on ice to a biosafety hood. The tumors were minced using sterilized scalpels and allowed to dissociate for 20-30 minutes in a 37°C water bath. Cell culture media was prepared using 2x the regular concentration of penicillin-streptomycin, and TrypLE was neutralized using an equivalent volume of this cell culture media. The mixture was filtered through a cell strainer tube (Corning; cat. no. 352235) and plated onto a tissue culture treated 10 cm dish, with cell culture media added to a total of 10 mL. The following day, upon cell adherence to the dish, the media was replaced. When cells reached 70-80% confluence, they were frozen down in media containing 2x the regular concentration of FBS and 10% DMSO. Cell pellets were simultaneously taken for downstream analyses, including TaqMan copy number assays and SMASH karyotyping. For subsequent phenotypic characterization, a fresh vial of cells was thawed and used in soft agar assays as previously described.

#### KRAS genotyping

The sequence of KRAS exon 2 was obtained from GRCh38 using Benchling. Primers were designed to amplify the genomic region surrounding exon 2 (Table S8). PCR was performed with SeqAmp DNA Polymerase (Takara Bio; cat. no. 63850), and PCR products purified and concentrated with the QIAquick PCR Purification kit (Qiagen; cat. no. 28106). PCR products were sent for Sanger sequencing with both forward and reverse primers to Azenta Biosciences. The ratio of wild type to mutant base (G to A) to assess allelic ratio of KRAS^WT^ to KRAS^G13D^ was assessed using EditR^114^.

#### Cell cycle analysis

One day prior to cell cycle analysis, 1.5-2 million cells were plated onto a 10 cm dish. Cells were harvested by trypsinization with TrypLE, followed by centrifugation at 1,000 rpm for 5 minutes, aspiration of the media, and resuspension of the cell pellet in 1 mL PBS. Cells were added dropwise to 4mL of ice cold 100% ethanol and fixed at -20°C for 5-15 minutes. Fixed cells were pelleted by centrifugation at 1,000 rpm for 5 minutes, resuspended in PBS containing 0.05% Triton X-100, 10 μg/mL RNAse A (Invitrogen, cat no. A32078), and 20 μg/mL propidium iodide (Life Technologies, cat. no. P3566), filtered through a cell strainer FACS tube cap. Cells were incubated for 30 minutes at room temperature in the dark. Cell cycle analysis was measured by flow cytometry (Miltenyi Biotech).

#### Senescence assay

Senescence-associated β-galactosidase staining was performed using a senescence β-galactosidase staining kit (Cell Signaling Technology, cat. no. 9860) according to the manufacturer’s instructions. Briefly, in a 6-well plate, the medium was discarded, and the cells were rinsed once in PBS. The cells were then fixed in a fixative reagent for 15 minutes at room temperature. The cells were then rinsed twice in PBS and incubated overnight at 37°C without CO2 in the β-galactosidase staining solution. The cells were imaged and analyzed under a light microscope (20X magnification, Olympus CKX53).

#### CRISPRi competition assays

CRISPRi competition assays were performed as described in ref^61^. In brief, A2780 cells were transduced with a dCas9-KRAB construct (Addgene #85969). dCas9-KRAB expressing TagBFP+ cells were selected for through bulk sorting, and dCas9-KRAB expression was confirmed through transduction of MCM2 guides with known biological activity. For competition assays, dCas9-KRAB expressing cells were transduced with guides in vectors co-expressing mCherry that targeted candidate genes, as well as negative control guides targeting the non-coding AAVS1 locus and positive control guides targeting the essential replication gene MCM2. Three days post-transduction, cells were subject to flow cytometry analysis (Miltenyi Biotech) to assess the starting percentage of mCherry+ cells. Cells were re-plated, and timepoints were taken every three days through the conclusion of the assay at the fifth timepoint. To normalize for differences in starting percentages of mCherry+ cells, fold change, defined as (starting % positive)/(% positive at timepoint), was calculated at each timepoint.

#### Assessing cellular sensitivity to UCK2 poisons and other chemotherapeutic agents

A 72-hour drug assay was performed to assess cellular sensitivity to the nucleotide analogs RX-3117 (MedChemExpress; cat. no. HY-15228) and 3-deazauridine (Cayman Chemical; cat. no. 23125) in both A2780, A2780 UCK2-overexpression, and MCF10A cell lines. One day prior to the addition of the nucleotide analogs, 25,000 cells were plated into individual wells of a 6-well plate. The following day, upon cell adherence to the plate, the media was discarded and replaced with drug-containing media at varying concentrations. The cells were grown in the presence of the drug for 72 hours. Following this, the cells were harvested and counted. A similar protocol was followed to assess the sensitivity of A2780 cells to doxorubicin (Selleck Chemicals; cat. no. S1208), gemcitabine (Selleck Chemicals; cat. no. S1714), and olaparib (Selleck; cat. no. S1060).

### OTHER GENETIC MANIPULATIONS

#### Generation of control clones

##### Rosa26 gRNA clones

A CRISPR guide was designed to target the noncoding Rosa26 locus and cloned into the Lenti-Cas9-gRNA-GFP vector (Addgene #124770) as described above. Cells were transfected with the Rosa26 targeting gRNA CRISPR plasmid using Lipofectamine 3000, and GFP+ cells were single cell sorted onto 96-well plates. Clonal cell lines were established and subject to karyotypic validation through SMASH karyotyping prior to phenotypic characterization.

##### 1q, 7p, and 8q gRNA clones

Cells were transfected with centromere-targeting gRNA CRISPR plasmids using Lipofectamine 3000 and clonal cell lines were established as described above for ReDACT-CO (Table S5A). Clones that maintained an extra copy of the targeted chromosome, as revealed by TaqMan copy number assays and SMASH-seq, were used as control clones.

##### Olfactory gene deletions

CRISPR guides were designed to flank the coding sequence of the targeted olfactory gene and cloned into the Lenti-Cas9-gRNA-GFP vector (Addgene #124770) as described above. Cells were transfected with the olfactory gene deletion guides using Lipofectamine 3000, and single GFP+ cells were sorted onto 96-well plates. Clonal cell lines were established and screened for single copy deletions of the targeted olfactory genes with TaqMan copy number assays as described above. To exclude the possibility of a chromosome truncation event, additional TaqMan copy number assays were performed with probes telomeric to the targeted gene. Clones with single copy deletions of the targeted olfactory gene were subject to karyotypic validation through SMASH-seq prior to phenotypic characterization.

##### Generation of Cassette Deletion Clones

A2780 and AGS integrant clones were co-transfected with a centromere targeting gRNA and a cassette proximal gRNA using Lipofectamine 3000. Cassette deletion was selected for with 10 µM ganciclovir (Sigma Aldrich, cat. no. G2536). Following ganciclovir selection, dsRed or GFP negative cells were single cell sorted onto 96-well plates, and clonal cell lines were established. Clones were screened for cassette loss with PCR primers spanning the deletion region (Table S8) and cassette deletion independent of any other karyotypic alterations was confirmed through SMASH karyotyping.

#### Deletion of a single copy of MDM4

CRISPR guides were designed to flank the coding sequence of the targeted gene and cloned into the Lenti- Cas9-gRNA-GFP vector (Addgene #124770) as described above. For dual guide plasmids, dual guide gene blocks were ordered from IDT and cloned into the Lenti-Cas9-gRNA-GFP vector using NEBuilder HiFi DNA Assembly (New England Biolabs, cat. no. E2621L). CRISPR gRNA sequences are listed in Table S5A. Cells were transfected with the gene deletion guides using Lipofectamine 3000, and GFP+ cells were single cell sorted onto 96-well plates. Clonal cell lines were established and screened for single copy deletions of targeted genes with TaqMan copy number assays as described above. To exclude the possibility of a chromosome truncation event, additional TaqMan copy number assays were performed with probes telomeric to the targeted gene. Clones with single copy deletions of the targeted genes were subject to karyotypic validation through SMASH karyotyping prior to phenotypic characterization.

#### Transformation of MCF10A with HRAS^G12V^

pBABE-HRASV12-Hyg (Addgene #195143) was generated by cloning hRASV12 from pBABE-HRASV12-puro (Addgene #9051) into pBABE-hyg (Addgene #1765) by digestion of both plasmids with BamHI and SalI, gel purification of digested DNA, ligation, and transformation into Stbl3 E. coli. The plasmid sequence was verified by sequencing. Viral preparation and transduction were performed as described above.

#### Derivation of TP53-knockout clones

TP53-targeting gRNAs were designed and cloned into the Lenti-Cas9-gRNA-GFP vector (Addgene #124770) for generating TP53-KO clones. In parallel, an AAVS1-targeting gRNA was utilized to generate isogenic TP53- WT control clones. A2780 cells were transfected with these CRISPR gRNA plasmids and GFP+ cells were single cell sorted onto 96-well plates. Clonal cell lines were established and screened for TP53-KO through western blotting. Clones were subject to karyotypic validation through SMASH karyotyping prior to phenotypic characterization.

#### Generation of HAP1 UCK2-KO cells

HAP1 cells harboring a CRISPR-induced frameshift mutation in the UCK2 gene were purchased from Horizon Discovery (cat. no. HZGHC007067c006). Loss of UCK2 expression was verified by western blotting.

#### Generation of UCK2-overexpressing cells

A2780 cells were transduced with pLV-Bsd-CMV-hUCK (Addgene #195141) and selected with 4µg/mL blasticidin (InvivoGen; cat. no. ant-bl-1). The plasmid sequence was verified by whole-plasmid sequencing.

### COMPUTATIONAL APPROACHES

#### Copy number timing analysis

Raw whole genome sequencing (WGS) data in Bam or Fastq formats were downloaded from public databases provided by the original publications^28, 29, 115^. We used ith.Variant pipeline (https://github.com/SunPathLab/ith.Variant) to call somatic copy number alterations (SCNA) and point mutations and to determine the clonality of these somatic variants^116^. In total, 38 Breast cancer samples (BRCA) from 21 patients and 37 Melanoma (MEL) primary tumor samples (all paired with normal control samples) passed our WGS data quality control and were included in the analysis. To reconstruct the evolutionary history of SCNAs in patient tumors, we applied BUTTE^117^ to infer the initiation time of clonal CN gains. We define the initiation time as the time fraction when the first gain occurs for a clonal SCNA. In brief, BUTTE estimates the initiation time of complex gains (or gains involving multiple steps) by modeling the quantitative relationship between point mutations and paths of copy number events. To do so, it first adopts the expectation–maximization (EM) algorithm to find the allele state distribution of point mutations. BUTTE then either directly solves the timing for SCNAs with identifiable CN history matrices^118^ or adopts linear programming to calculate the upper bounds of the initiation time if the underlying linear system is underdetermined (e.g., Multiple history matrices exist for an SCNA)^117^. We identified clustered gains by clustering the inferred timing via nonparametric density estimation^119^. Genome doubling was identified as the cluster containing more than 40% of the segments.

#### Detecting recurrent early gains

To identify genomic regions exhibiting recurrent early gains across patients, we scanned hg38 genome with bins of 1 million bp in size and ranked the bins in each sample according to the timing of respective initiating gain. The timing values are jittered to avoid ties. We then subtract from each bin the middle rank of the respective sample. The middle rank is the expectation value of the ranks under the assumption that the null hypothesis holds: no regions show recurrent early gain across patients. For each tumor type, we summed the resulting ranks across patients for each bin. Division of the rank sums by its standard deviation yields normalized rank sums which approximately follow a standard normal distribution if the null hypothesis is fulfilled. A large negative normalized rank sum would reject the null hypothesis and indicate recurrent early initiating gains. To evaluate the frequency of gains for each genomic bin across patients, we ranked the segment mean (read depth ratio between the tumor and normal samples) as we did for the timing values, and then performed the same rank sum normalization. A large positive normalized rank sum for the segment mean would suggest frequent gains across patients.

#### Analysis of mutual exclusivity between aneuploidy and oncogenic mutations

To analyze the mutual exclusivity between aneuploidy and oncogenic mutation, we started from the MSK-MET^34^ dataset available via the public cBioPortal datahub^120^. In order for a given chromosomal arm and oncogene to be compared, we required that the data pass a set of quality filters. We required all cancer types to have ≥ 25 patients and selected a representative sample for each patient based on available mutation and arm gain data. We built sets of genes and arms by cancer type by applying additional requirements by cancer. The sets of genes we analyzed started from the IMPACT-505 geneset and we required each gene be mutated in ≥ 2% of patients by cancer type. The sets of arms we analyzed started from all previously generated arm gain data by sample and each arm was required to exceed a minimum of 100 patients or ≥ 2% of patients experiencing arm gain by cancer type. Once we had our cleaned sets of genes and arms by cancer, we calculated contingency tables for each combination and conducted a two-sided Fisher’s exact test to compute p-values and odds ratios for each gene X arm combination. We converted these p-values into Z scores for further comparisons and applied the sign of the negative log2 of the odds ratio to represent the direction of the relationship; therefore, a large positive Z would suggest significant mutual exclusivity, while a large negative Z would suggest significant co-occurrence. We repeated this process in a pancancer analysis by utilizing the same cleaned data and pooling all patients, genes, and arms into a single cohort. All mutual exclusivity analysis was automated using Nextflow^98^ and Conda^121^, starting from downloading MSK-MET data through to generation of Figures 1D-E and S1.

#### Analysis of copy number alterations and patient survival

To examine the relationship between copy number gains, mutations, and cancer patient outcomes in TCGA, univariate Cox proportional hazards models were used as described in refs^122, 123^. In brief, TCGA copy number data (broad.mit.edu_PANCAN_Genome_Wide_SNP_6_whitelisted.seg, available at https://gdc.cancer.gov/ about-data/publications/pancanatlas) and TCGA mutation data (mc3.v0.2.8.PUBLIC.maf.gz, available at https://gdc.cancer.gov/about-data/publications/pancanatlas) were combined with patient outcomes data described in ref^124^. Selection of the clinical endpoint for each cancer type was based on the recommendations provided by ref^124^ based on data quality, cohort size, and the number of events that were observed. TCGA copy number data was generated as relative copy number values for particular chromosomal intervals. This data was translated to produce a single copy number value on a per-gene basis, based on the observed copy number at each gene’s transcription start site. This annotation was performed using mapping data from GENCODE v32^125^. Z scores for copy number changes and for mutations in common cancer driver genes were calculated using Cox proportional hazards regression. For the copy number analysis presented in Figure S2, genes were mapped back to chromosome bands, and the top-scoring gene (based on the pan-cancer Stouffer’s Z score) was identified for each band. For the mutation analysis presented in Figure S2, a gene was considered to be mutated if there was a single non-synonymous mutation at any codon within the gene. Non-synonymous mutations included: missense, nonsense, frameshift deletion, splice site, frameshift insertion, inframe deletion, translation start site, nonstop mutation, and in-frame insertion. Additional clinical data presented in Figure S2E were downloaded from cBioportal^120^.

#### Analysis of aneuploidy-associated gene expression

Chromosome copy number data for cancers from the TCGA was downloaded from ref^1^. Chromosome copy number data for the Cancer Cell Line Encyclopedia was downloaded from ref^93^. Gene expression data for cancers from the TCGA was downloaded from the TCGA PanCanAtlas (EBPlusPlusAdjustPANCAN_ IlluminaHiSeq_RNASeqV2.geneExp.tsv, available at https://gdc.cancer.gov/about-data/publications/ pancanatlas). Gene expression data for the Cancer Cell Line Encyclopedia was downloaded from DepMap (www.depmap.org)^126^.

### DATA VISUALIZATION

Scientific illustrations were generated with Biorender. Most graphs were generated using GraphPad Prism. Boxplots display the 25th, 50th, and 75th percentiles of colonies per field, while the whiskers represent the 10th and 90th percentiles. Unless otherwise indicated, bar graphs and XY plots display the mean ± SEM.

### DATA AND CODE AVAILABILITY

The code used to perform the TCGA survival analysis is available at https://github.com/joan-smith/comprehensive-tcga-survival. The code used to perform the mutual exclusivity analysis is available at github.com/shel tzer-lab/aneuploidy-addictions. The mass spectrometry data have been all deposited to the ProteomeXchange Consortium via the PRIDE PXD037956^127^. (To review the dataset please go to https://www.ebi.ac.uk/pride/login, and use the following login details: **Username:**; **Password: NavbYhDW**). RNA-Seq data has been deposited at GSE222379.

## SUPPLEMENTAL TABLES

**Table S1. Mutual exclusivity between chromosome gain events and mutations in cancer-associated genes.**

**Table S2. Association between chromosome copy number gains and cancer patient outcome.**

**Table S3. Aneuploidy-loss cell lines generated in this work.**

**Table S4. Sources of the cell lines used in this work.**

**Table S5. CRISPR and CRISPRi gRNA sequences used in this work.**

**Table S6. TaqMan copy number probes used in this work.**

**Table S7. TaqMan gene expression probes used in this work.**

**Table S8. Primers and oligonucleotides used in this work.**

## Supplemental Text 1. Considering alternate explanations for the loss of fitness in engineered 1q-disomic cancer cells

In order to generate isogenic cancer cells that have or lack specific aneuploid chromosomes, we developed and applied a suite of CRISPR tools for chromosome engineering called ReDACT. We discovered that eliminating the trisomy of chromosome 1q severely compromised malignant potential in multiple independent cancer cell lines. We considered and rigorously evaluated the possibility that the loss of fitness observed among the 1q- disomic clones could be a consequence of our chromosome engineering methodologies, rather than the subsequent change in cellular karyotype. However, multiple lines of evidence indicate that this loss of fitness is best explained as a specific outcome of eliminating trisomy-1q and not a consequence of our experimental approach:

### The use of CRISPR

All three ReDACT techniques that we applied utilized CRISPR to induce aneuploidy-loss events. In order to assess whether CRISPR itself could compromise malignant growth to the degree that we observed upon elimination of the 1q trisomy, we generated and tested a set of 28 control clones that were subject to various CRISPR manipulations. These clones include:

1) Cell lines harboring a CRISPR-mediated integration of the HSV-TK cassette that were not treated with ganciclovir to select for 1q-loss,
2) Cell lines in which the HSV-TK cassette was deleted by transfecting cells with two gRNAs targeting immediately upstream and downstream of the integrant coupled with ganciclovir selection, which resulted in a segmental deletion of the cassette while leaving the rest of 1q unaffected,
3) Cell lines subjected to CRISPR-mediated cutting with a 1q-targeting gRNA, in which the lesion was repaired without causing chromosome loss,
4) Cell lines subjected to CRISPR-mediated cutting with a gRNA targeting the non-coding Rosa26 locus,
5) Cell lines in which dual CRISPR guides were used to generate segmental deletions on chromosome 1q of a gene encoding a non-expressed olfactory receptor,
6) Cell lines in which CRISPR was used to delete a terminal segment on chromosome 1q, eliminating the telomere and decreasing the copy number of 26 out of 968 protein-coding genes on the chromosome.

Every control clone that we tested exhibited significantly better anchorage-independent growth compared to the 1q-disomic clones that we derived (Fig. S8). Additionally, we note that the control clones generated by the segmental deletion of the HSV-TK gene were subjected to three independent CRISPR-induced DNA breaks (one to integrate HSV-TK and then two to produce the segmental deletion), which is more breaks than all 1q- disomic clones were subjected to.

To further verify that the phenotypes observed upon elimination of the 1q-trisomy are specifically a result of that karyotype alteration, we applied ReDACT-CO to eliminate trisomies of 1q, 7p, and 8q from the same cell line (Fig. 3A). If the use of ReDACT-CO is the cause of the reduced fitness upon 1q-loss, then we would expect that all aneuploidy-loss clones would be impaired to a similar degree. However, we observed that loss of the 7p or 8q trisomies resulted in a significantly milder phenotype compared to the effects of 1q-loss (Fig. 3C-D).

Finally, if an off-target effect of CRISPR is the cause of the reduced fitness upon 1q-loss, then we would not expect to see any selective pressure to restore the 1q trisomy. However, upon prolonged growth of the 1q- disomic clones *in vitro* or *in vivo*, we observed that many cell populations spontaneously recover an extra copy of chromosome 1q, and these 1q-restored cells exhibit improved colony-formation ability relative to the 1q- disomic clones (Fig. 4A-F). In total, these assays provide multiple independent lines of evidence that the reduced fitness of the 1q-disomic clones cannot be attributed solely to the effects of CRISPR.

### Ganciclovir selection

In the ReDACT-NS approach, several 1q-disomic clones were generated by integrating the HSV-TK gene onto chromosome 1q and then selecting for aneuploidy-elimination via treatment with ganciclovir (Fig. 2A). As a specific control for this protocol, we also generated a series of clones in which the HSV-TK-expressing parental cells were transfected with two gRNAs that cut immediately upstream and downstream of the HSV-TK cassette, and then the cells were treated with ganciclovir (Fig. S8A). These clones acquired ganciclovir resistance due to a segmental deletion of the HSV-TK gene, rather than loss of the entire chromosome arm. We subsequently observed that these ganciclovir-resistant control clones exhibited consistently superior anchorage-independent growth compared to clones that had lost the 1q-trisomy (Fig. S8D-E). Additionally, we note that two of our aneuploidy-elimination methods – ReDACT-TR and ReDACT-CO – do not utilize ganciclovir selection, and the phenotypes that we observed across independent 1q-disomic clones were similar regardless of the methods applied to generate them. In total, these findings suggest that any detrimental effects of ganciclovir selection are unable to fully account for the loss of fitness observed in the 1q-disomic clones.

### Loss of telomere protection

In the ReDACT-NS and ReDACT-CO approaches, our aneuploidy elimination techniques may result in the loss of telomere protection on the targeted chromosome arm. We therefore investigated whether the loss of telomere protection could be sufficient to explain the phenotypes of our 1q-disomic clones. First, we transfected cells with a gRNA targeting a subtelomeric region on chromosome 1q and we isolated a control clone harboring a terminal chromosomal truncation. This clone maintained the ability to grow under anchorage-independent conditions at wild-type levels (Fig. S8C). Second, if loss of telomere protection on a single chromosome arm is sufficient to inhibit malignant potential, then we would expect this phenotype to be consistent across different chromosomes. However, we applied ReDACT-CO to eliminate the trisomies of chromosome 7p and 8q from A2058 cells, and we observed that the 7p-disomic and 8q-disomic clones exhibited consistently superior fitness compared to 1q- disomic clones obtained using the same techniques in the same cell line (Table S3). Third, a subset of our 1q- disomic clones were created using ReDACT-TR, in which the CRISPR-induced DNA break was repaired with an artificial telomere. As noted above, the phenotypes that we observed across independent 1q-disomic clones were similar regardless of the methods applied to generate them (Fig. 2). In total, these findings suggest that the loss of telomere protection on a single chromosome arm is unable to fully account for the compromised fitness observed in the 1q-disomic clones.

## Supplemental Text 2. Considering the loss of specific point mutations on chromosome 1q as an explanation for the loss of fitness in engineered 1q-disomic cancer cells

Deletion of a chromosome not only decreases the dosage of any genes encoded on the targeted chromosome, it may also cause the loss of any point mutations encoded on that chromosome. Correspondingly, we considered the possibility that the effects of 1q-loss could be mediated in part by eliminating unique driver mutations that these cell lines had acquired on chromosome 1q. To explore this possibility, we evaluated all non-synonymous mutations on chromosome 1q in the 1q-trisomic cancer cell lines used in this study. Using data acquired from DepMap, we found 25 1q mutations in A2780, 19 1q mutations in A2058, and 52 1q mutations in AGS. We cross- referenced these mutations with the Catalogue of Somatic Mutations in Cancer (COSMIC) database to examine if any mutations were recurrently observed or causally implicated in human cancers. None of the 96 mutations present on chromosome 1q in A2780, A2058, and AGS were identified as mutational hotspots in the COSMIC database, and none of the genes affected by mutations were included in the Cancer Gene Census. Next, we investigated the list of cancer driver genes identified by Vogelstein et al.^129^, and we found that none of the 1q genes affected by mutations in these cell lines were annotated as likely drivers. Lastly, we examined MSK- IMPACT, a panel of 505 genes associated with both common and rare cancers^130^. None of the 1q mutated genes are included in this panel. For these reasons, we believe that the mutations found on chromosome 1q in these cell lines likely represent passenger events, rather than cancer drivers. Nonetheless, we do not rule out the possibility that the loss of specific point mutations could influence the effects of aneuploidy-elimination in other experiments. For instance, as described in Figure 4G, we speculate that the effects of gaining chromosome 12 in HCT116 is mediated in part by the acquisition of an extra copy of the mutant KRAS^G13D^ allele, and loss of a chromosome containing mutant KRAS may have different consequences than loss of a chromosome containing wild-type KRAS.

---

**Figure S1.**
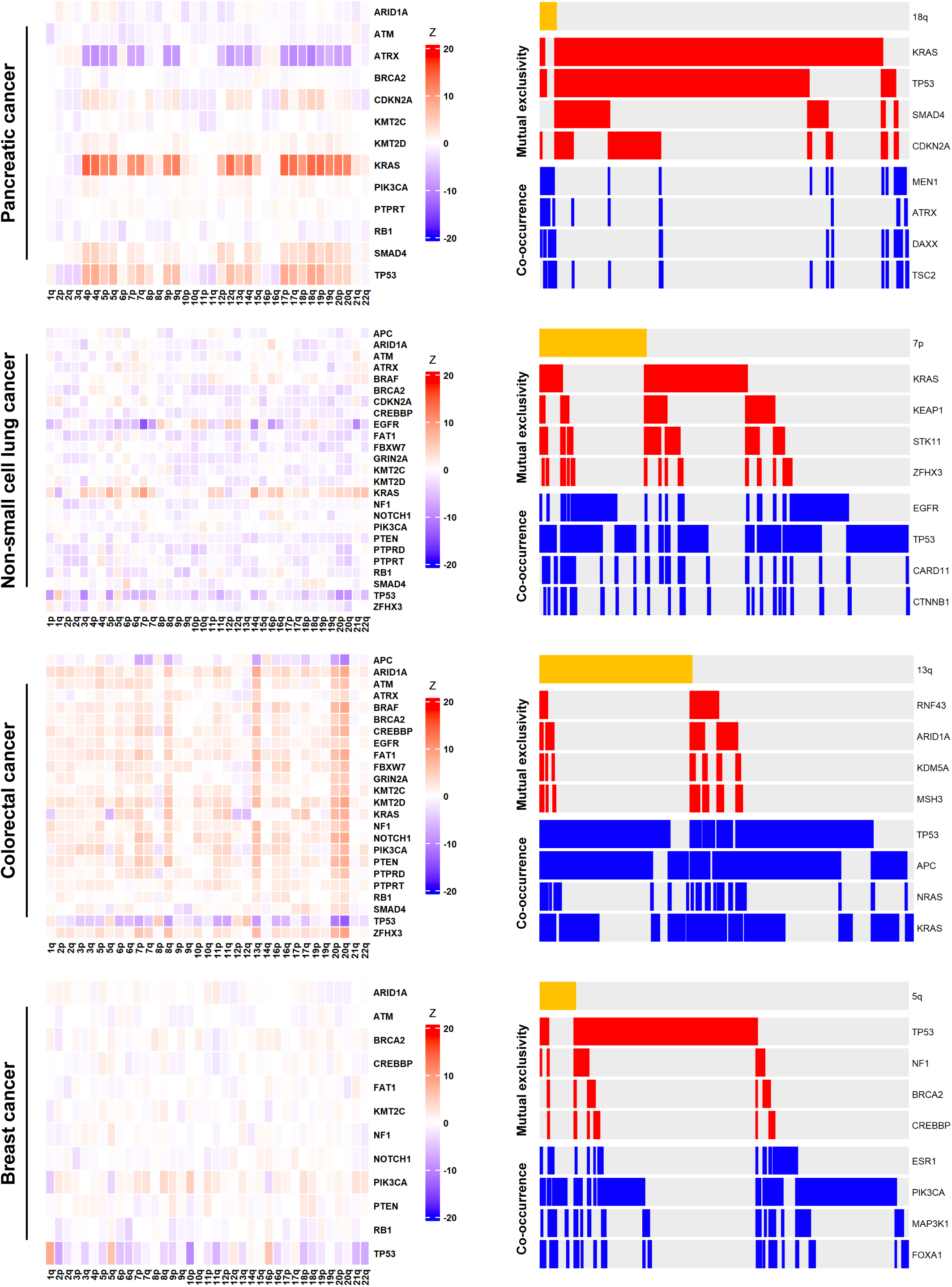
Mutual exclusivity between chromosome arm gains and mutations in cancer driver genes in individual cancer types. (Left) Heatmaps are displayed demonstrating mutual exclusivity (in red) and co-occurrence patterns (in blue) between chromosome arm gains and mutations in common cancer driver genes within four individual cancer types: pancreatic cancer, non-small cell lung cancer, colorectal cancer, and breast cancer. (Right) Oncoprint panels highlighting mutual exclusivity and co-occurrence patterns within individual cancer types for specific chromosome gain events. The complete results of this analysis are included in Table S1.

**Figure S2.**
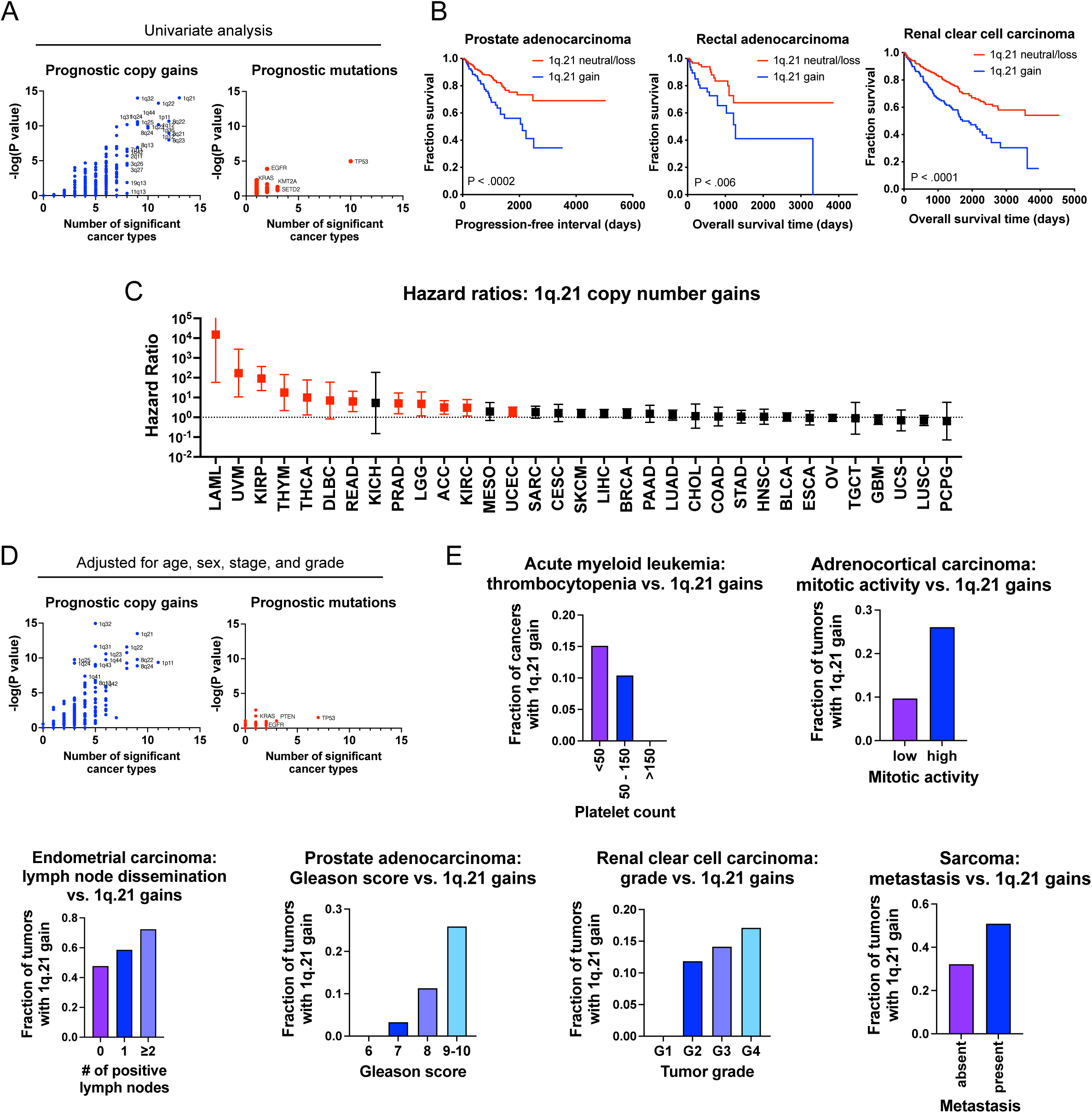
Specific copy number gains, particularly involving chromosome 1q.21, are associated with disease progression and patient death. (A) A scatterplot displaying the relationship between copy number gains for each chromosome band (left) and mutations (right) and patients outcomes across 32 cancer types from the TCGA^123^. Significance was determined by calculating univariate Cox proportional hazards regression models for each data type. The complete results for this analysis are presented in Table S2A. (B) Kaplan-Meier plots displaying the relationship between 1q.21 copy number gains and disease progression in prostate adenocarcinoma, rectal adenocarcinoma, and renal clear cell carcinoma. (C) A forest plot showing hazard ratios and 95% confidence intervals for Cox proportional hazards regression between 1q.21 copy number and patient outcome for each of the indicated cancer types. The hazard ratios plotted in red represent those that are significant at a p < 0.05 threshold. (D) A scatterplot displaying the relationship between copy number gains for each chromosome band (left) and mutations (right) and patients outcomes across 32 cancer types from the TCGA^123^. Significance was determined by calculating multivariate Cox proportional hazards regression models for each data type, adjusting each model for patient age and sex, and tumor stage and grade. The complete results for this analysis are presented in Table S2B. (E) Chromosome 1q.21 gains are associated with hallmarks of disease progression risk in various cancer types.

**Figure S3.**
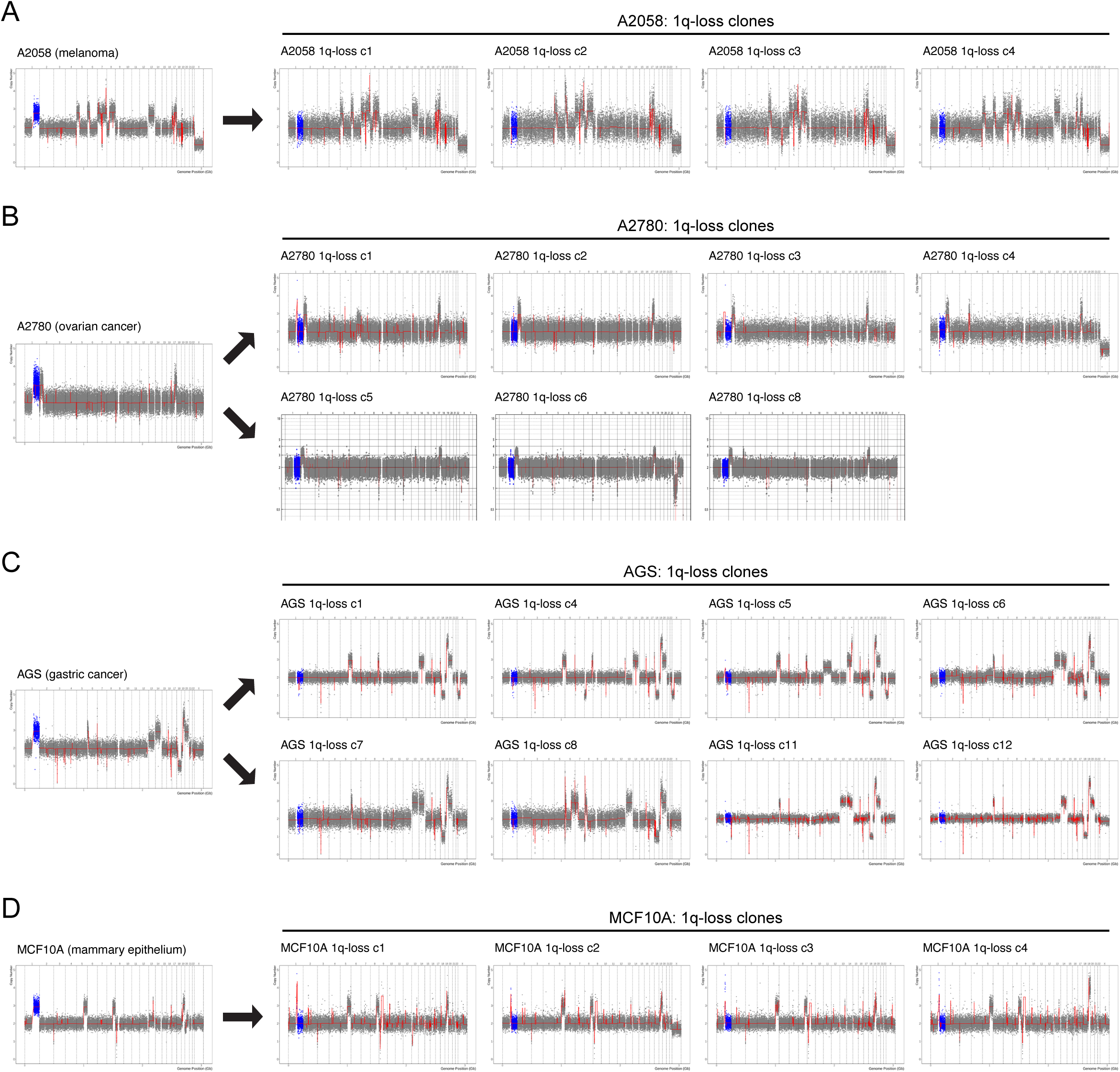
Additional karyotypes of 1q-disomic clones. (A) SMASH karyotypes of 1q-disomic clones generated from the 1q-trisomic cancer cell line A2058. (B) SMASH karyotypes of 1q-disomic clones generated from the 1q-trisomic cancer cell line A2780. (C) SMASH karyotypes of 1q-disomic clones generated from the 1q-trisomic cancer cell line AGS. (D) SMASH karyotypes of 1q-disomic clones generated from the 1q-trisomic immortalized mammary epithelial cell line MCF10A. For all karyotypes in A-D, chromosome 1q is highlighted in blue. A complete list of aneuploidy-loss clones is included in Table S3.

**Figure S4.**
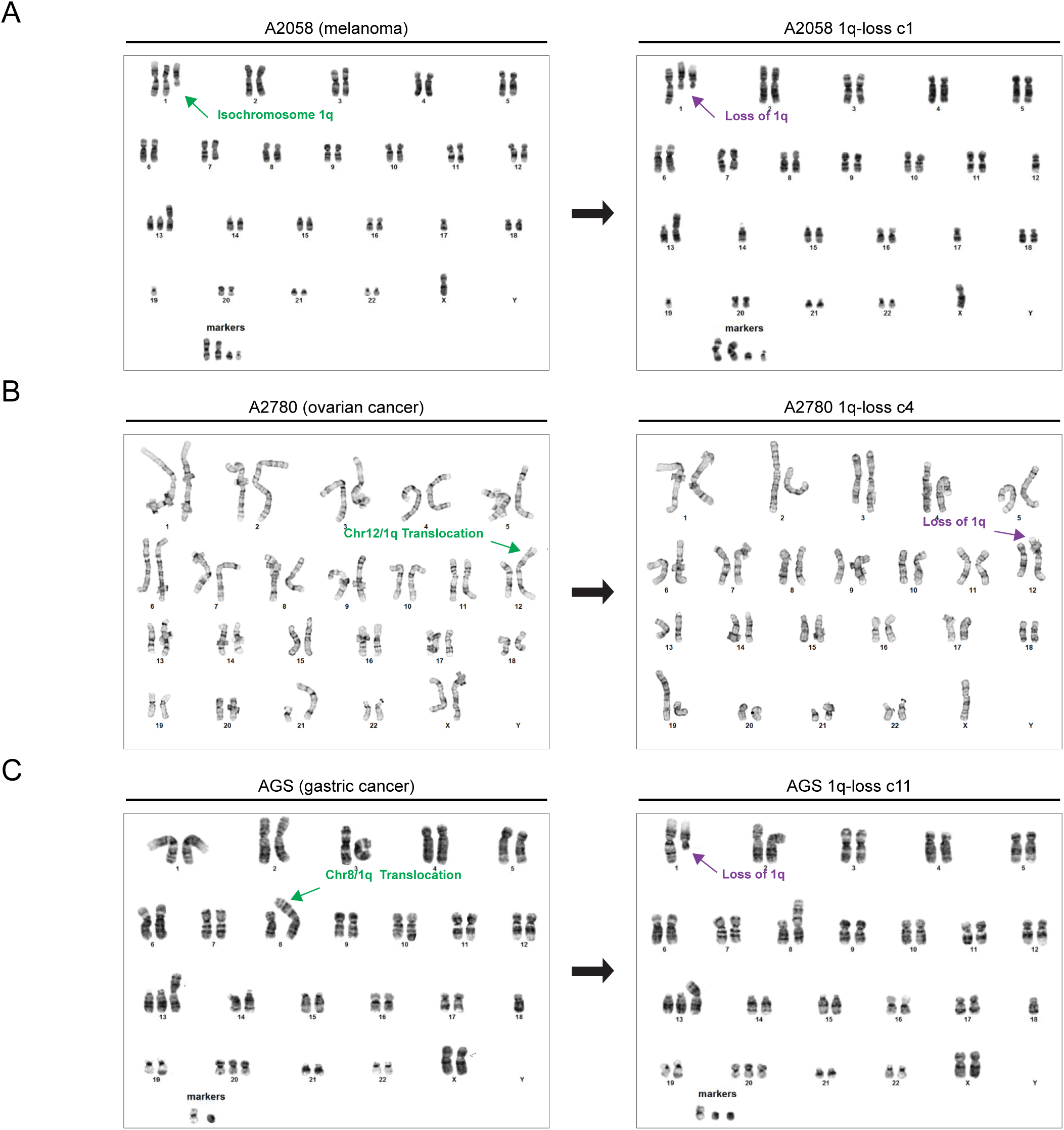
G-banded karyotypes of 1q-disomic clones. (A) G-banded karyotypes of A2058 and a 1q-disomic A2058 clone. The third copy of chromosome 1q in A2058 is present as an isochromosome [chr 1, iso(1q), del(1q)], and 1q-disomic clone has lost a copy of 1q from the intact chromosome 1. (B) G-banded karyotypes of A2780 and a 1q-disomic A2780 clone. The third copy of chromosome 1q in A2780 is translocated to chromosome 12q, and the 1q-disomic clone has lost this translocated copy of 1q. (C) G-banded karyotypes of AGS and a 1q-disomic AGS clone. The third copy of chromosome 1q in AGS is translocated to chromosome 8p, and the 1q-disomic clone has lost a copy of 1q from an intact chromosome 1. For all G-banded karyotypes in A-C, green arrows indicate the extra copy of chromosome 1q in parental cells and purple arrows indicate the deletion in the 1q-disomic clones.

**Figure S5.**
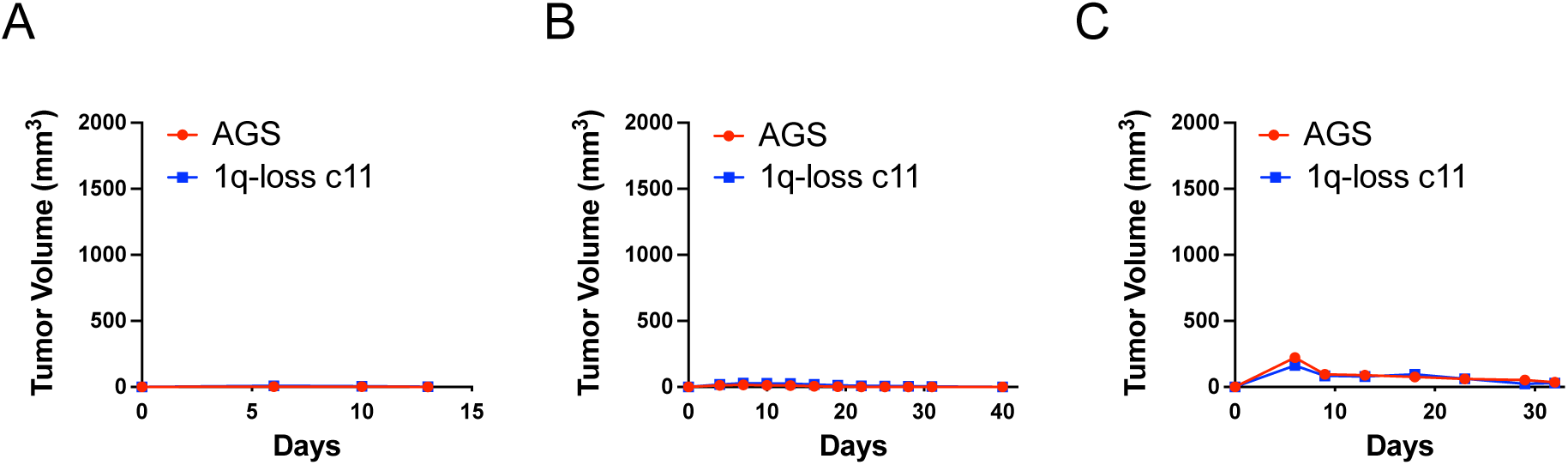
Wild-type AGS cells fail to grow as subcutaneous xenografts. (A) 5 million 1q-trisomic or 1q-disomic AGS cells were suspended in PBS and injected subcutaneously and contralaterally into 5-week-old female Nu/J mice. No tumor formation was observed. (B) 4 million 1q-trisomic or 1q-disomic AGS cells were suspended in PBS and injected subcutaneously and contralaterally into 6-week-old female NSG mice. No tumor formation was observed. (C) 15 million 1q-trisomic or 1q-disomic AGS cells were suspended in a 1:1 mixture of PBS and Matrigel and injected subcutaneously and contralaterally into 6-week-old female J:NU mice. No tumor formation was observed.

**Figure S6.**
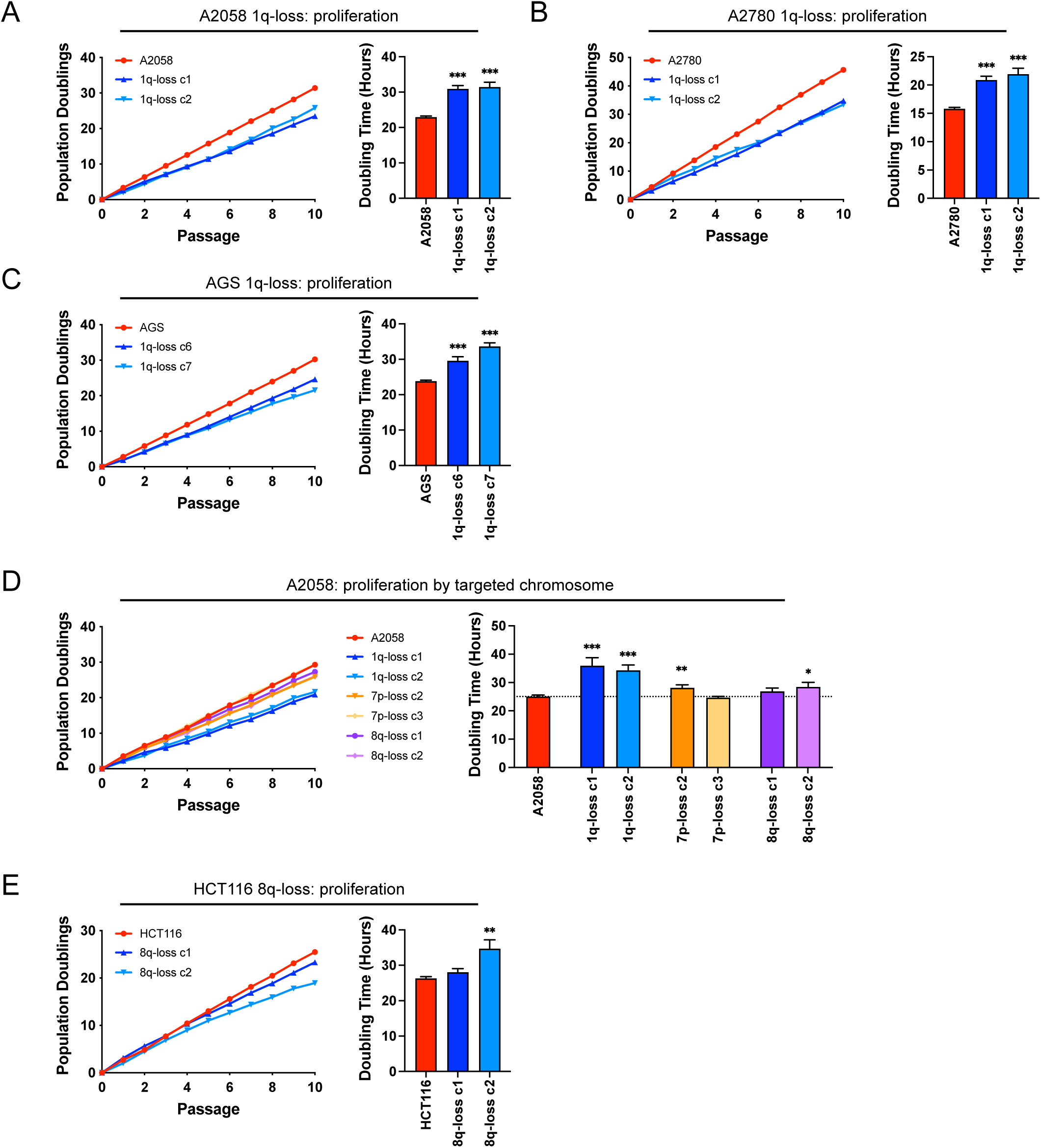
Elimination of trisomy-1q causes a moderate decrease in 2D cell proliferation. (A) Proliferation assays in 1q-trisomic and 1q-disomic A2058 cells. (B) Proliferation assays in 1q-trisomic and 1q-disomic A2780 cells. (C) Proliferation assays in 1q-trisomic and 1q-disomic AGS cells. (D) Proliferation assays in parental and 1q-disomic, 7p-disomic, or 8q-disomic A2058 cells. (E) Proliferation assays in 8q-trisomic and 8q-disomic HCT116 cells. For all proliferation assays in A-E, cumulative population doublings are shown on the left, and the doubling time calculated from these assays is shown on the right. Data are the mean ± SEM, n=10 passages. Data were analyzed by unpaired t-test. *p < 0.05, **p < 0.005, ***p < 0.0005

**Figure S7.**
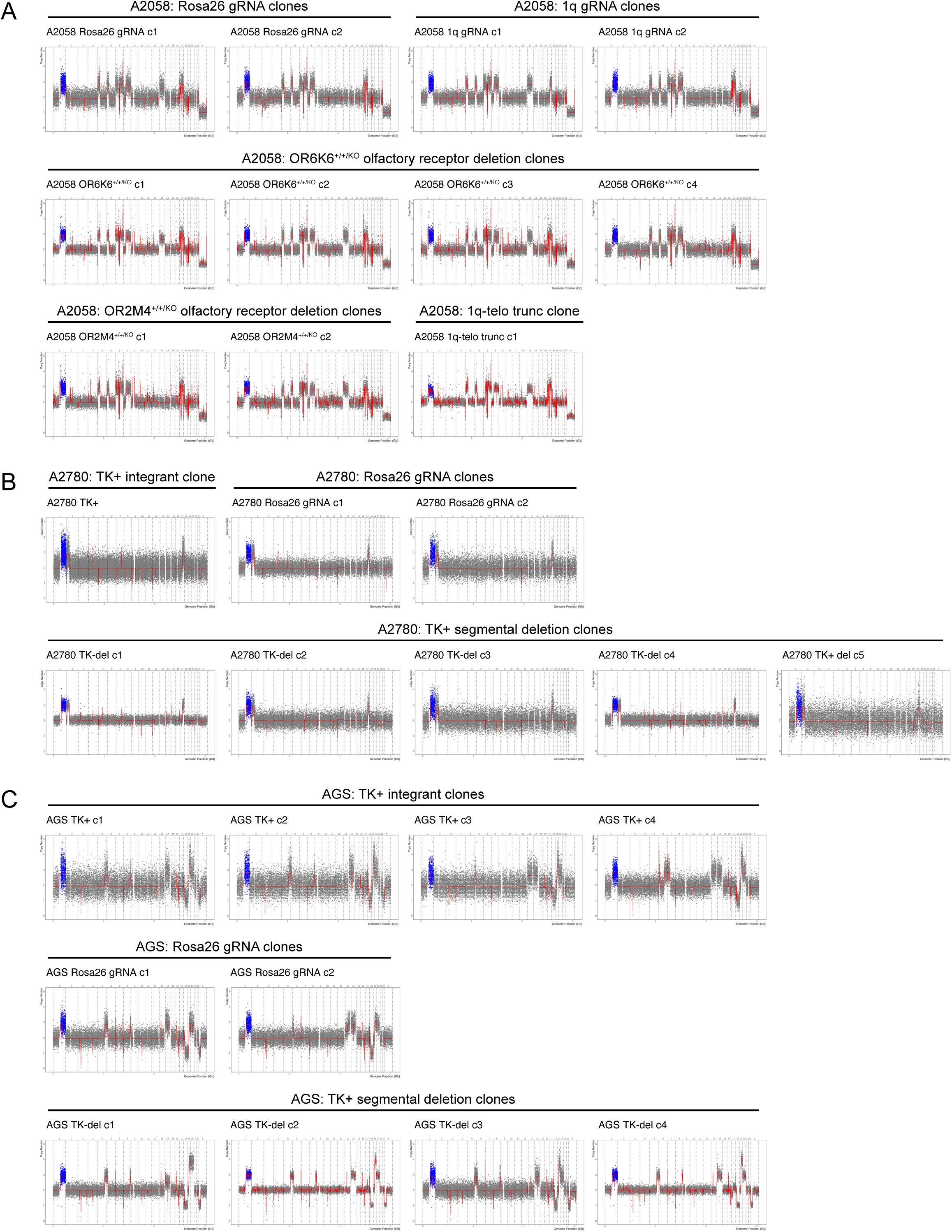
Karyotypes of control clones subjected to various CRISPR manipulations. (A) SMASH karyotypes of the control clones generated from the 1q trisomic cancer cell line A2058, with all clones maintaining 1q trisomy. (B) SMASH karyotypes of the control clones generated from the 1q trisomic cancer cell line A2780, with all clones maintaining 1q trisomy. (C) SMASH karyotypes of the control clones generated from the 1q trisomic cancer cell line AGS, with all clones maintaining 1q trisomy. For all karyotypes in A-C, chromosome 1q is highlighted in blue. Additional information on how these clones were generated is presented in Figure S8.

**Figure S8.**
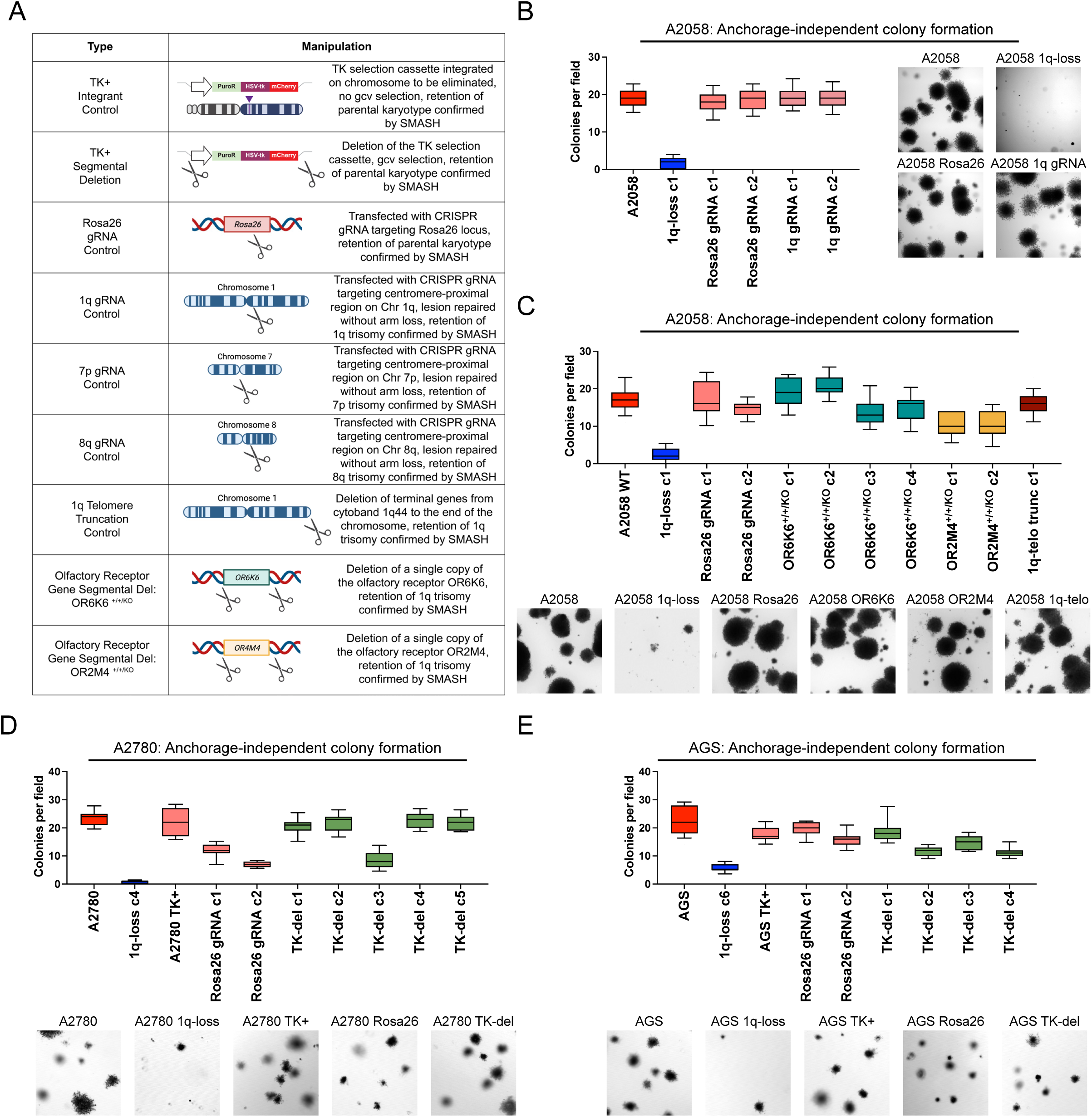
1q-disomic cancer cells exhibit uniformly worse growth in soft agar compared to control clones subjected to various CRISPR manipulations. (A) A schematic of the various strategies used to generate the control clones tested in this work. Note that the 7p and 8q gRNA control clones are tested in Figure S10. (B) A 1q-disomic clone exhibits worse anchorage-independent growth compared to Rosa26 gRNA and 1q gRNA control clones in A2058. (C) A 1q-disomic clone exhibits worse anchorage-independent growth compared to Rosa26 gRNA, olfactory receptor deletion, and 1q-telo truncation control clones in A2058. (D) A 1q-disomic clone exhibits worse anchorage-independent growth compared to TK+ integrant, Rosa26 gRNA, and TK-segmental deletion control clones in A2780. Note that these control clones display some variation in anchorage-independent growth, but every control clone grew better than a 1q-disomic clone. (E) A 1q-disomic clone exhibits worse anchorage-independent growth compared to TK+ integrant, Rosa26 gRNA, and TK-segmental deletion control clones in AGS. For anchorage-independent growth assays in B-D, boxes represent the 25th, 50th, and 75th percentiles of colonies per field, while the whiskers represent the 10th and 90th percentiles. Data are from n = 15 fields of view, and a representative trial is shown for each experiment.

**Figure S9.**
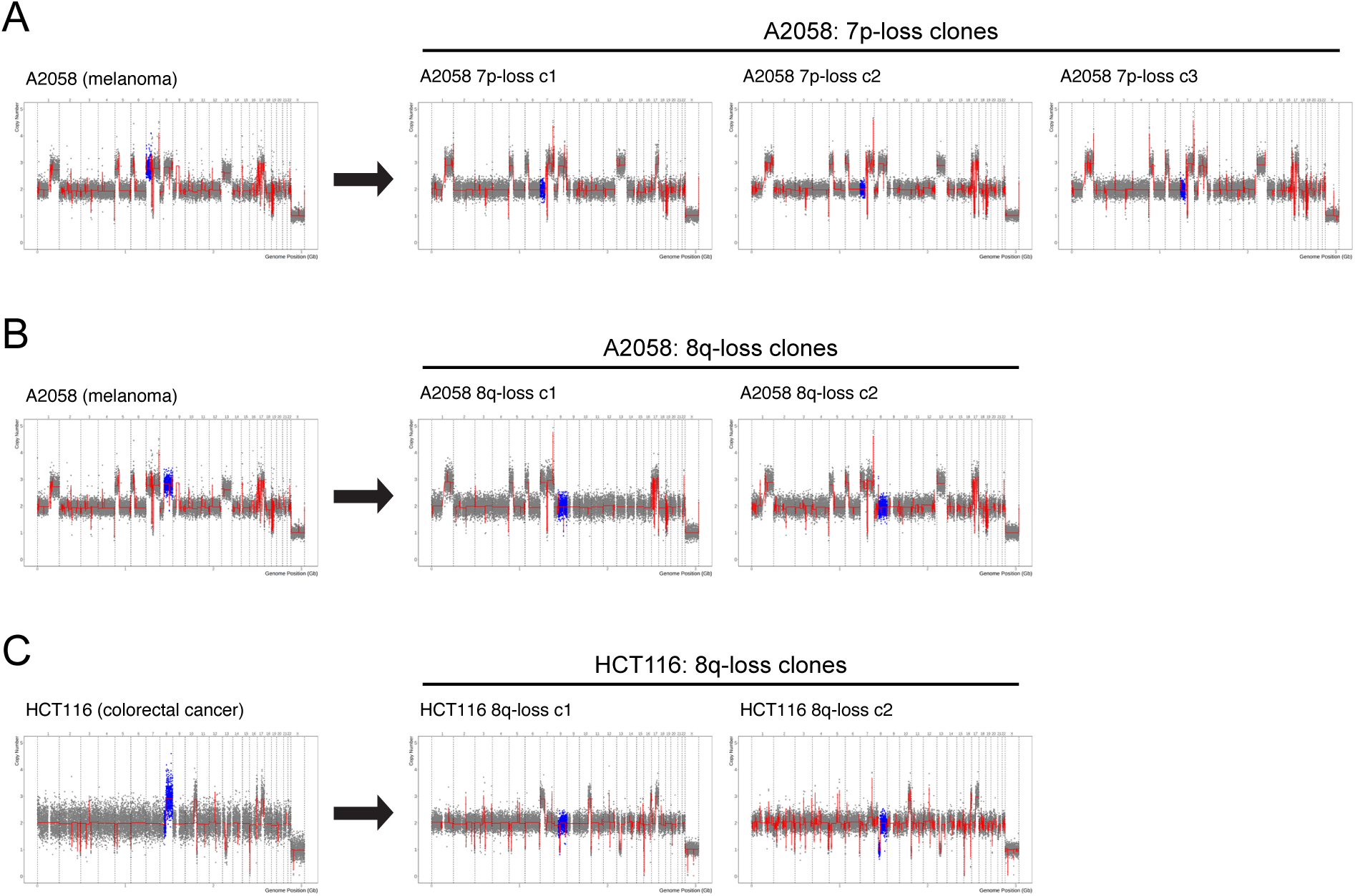
Additional karyotypes of 7p-disomic and 8q-disomic clones. (A) SMASH karyotypes of 7p-disomic clones generated from the 7p-trisomic cancer cell line A2058. Chromosome 7p is highlighted in blue. (B) SMASH karyotypes of 8q-disomic clones generated from the 8q-trisomic cancer cell line A2058. Chromosome 8q is highlighted in blue. (C) SMASH karyotypes of 8q-disomic clones generated from the 8q-trisomic cancer cell line HCT116. Chromosome 8q is highlighted in blue.

**Figure S10.**
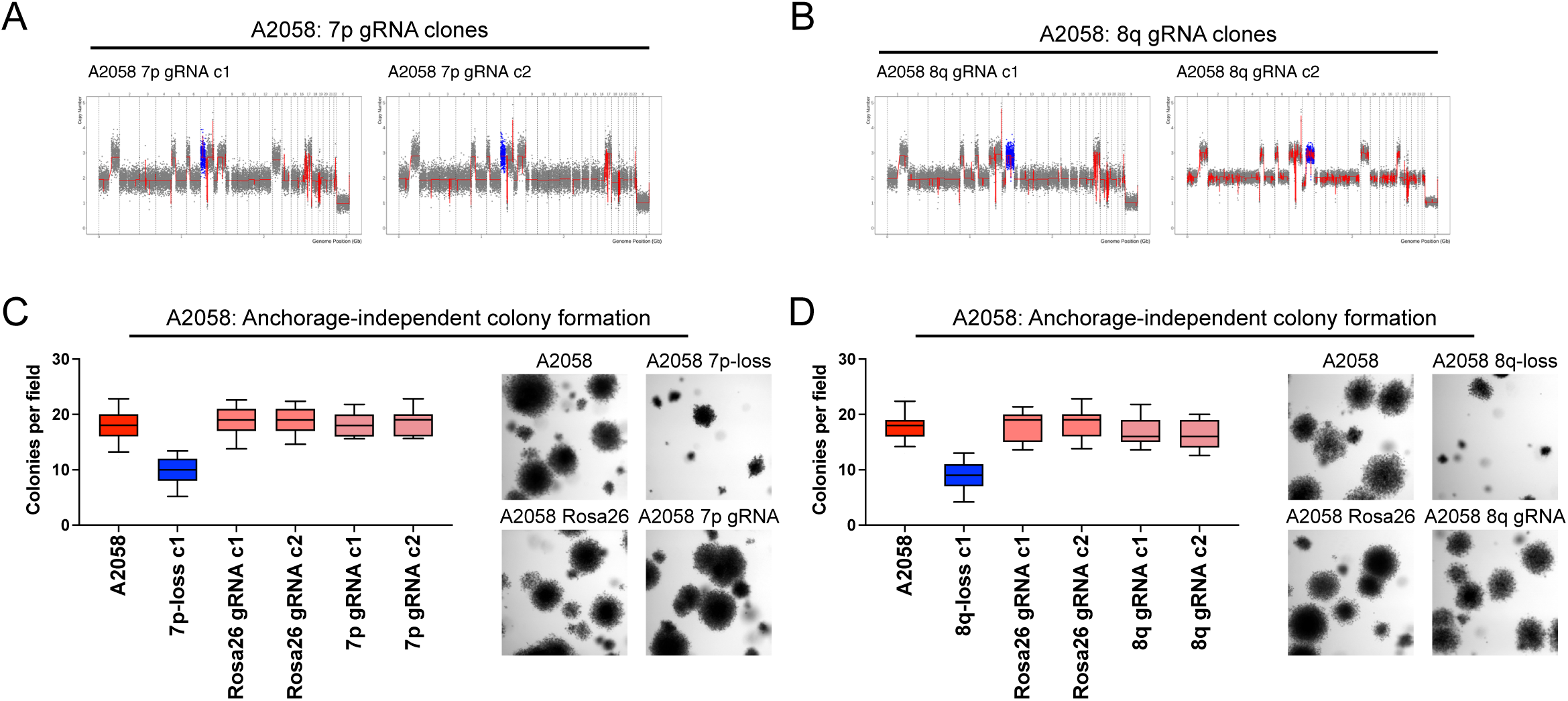
Karyotypes and anchorage-independent growth of A2058 7p and 8q control clones. (A) SMASH karyotypes of control clones generated by transfecting cells with the same CRISPR gRNA used to produce the 7p-disomic clones. Clones were selected that maintain the 7p trisomy. Chromosome 7p is highlighted in blue. (B) SMASH karyotypes of control clones generated by transfecting cells with the same CRISPR gRNA used to produce the 8q-disomic clones. Clones were selected that maintain the 8q trisomy. Chromosome 8q is highlighted in blue. (C) A 7p-disomic clone exhibits worse anchorage-independent growth compared to Rosa26 gRNA and 7p gRNA control clones in A2058. (D) An 8q-disomic clone exhibits worse anchorage-independent growth compared to Rosa26 gRNA and 8q gRNA control clones in A2058. For anchorage-independent growth assays in C-D, boxes represent the 25th, 50th, and 75th percentiles of colonies per field, while the whiskers represent the 10th and 90th percentiles. Unpaired t-test, n = 15 fields of view, data from representative trial. Representative images are shown next to each graph.

**Figure S11.**
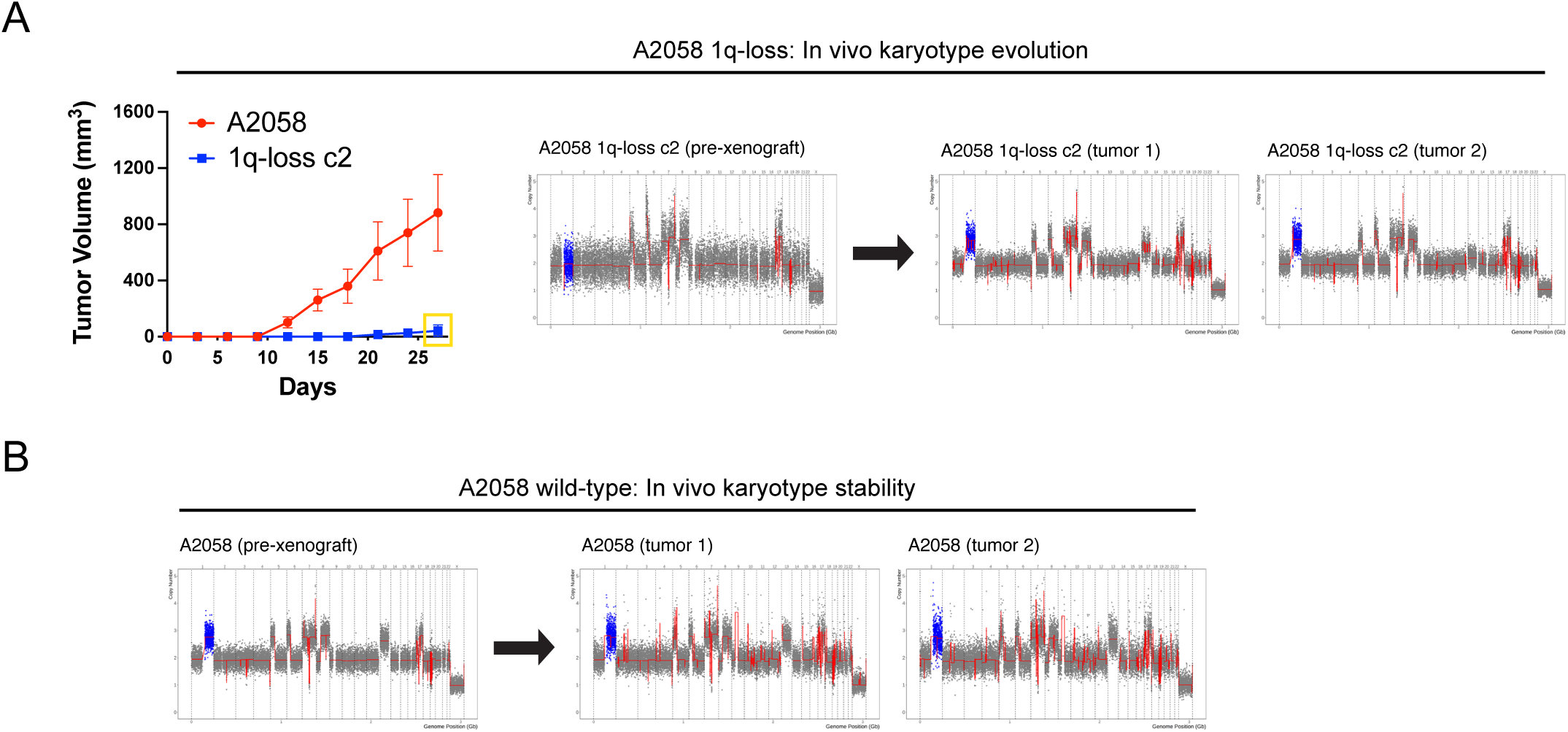
Karyotype analysis of additional 1q-trisomic and 1q-disomic A2058 xenografts. (A) 1q-disomic cells frequently evolve to recover a third copy of chromosome 1q during xenograft growth. (B) A2058 wildtype tumors exhibit karyotypic stability during xenograft growth. For the data shown in A-B, initial karyotypes for these lines prior to xenograft assays are shown on the left, and karyotypes of tumors following the xenograft assays are shown on the right. Chromosome 1q is highlighted in blue.

**Figure S12.**
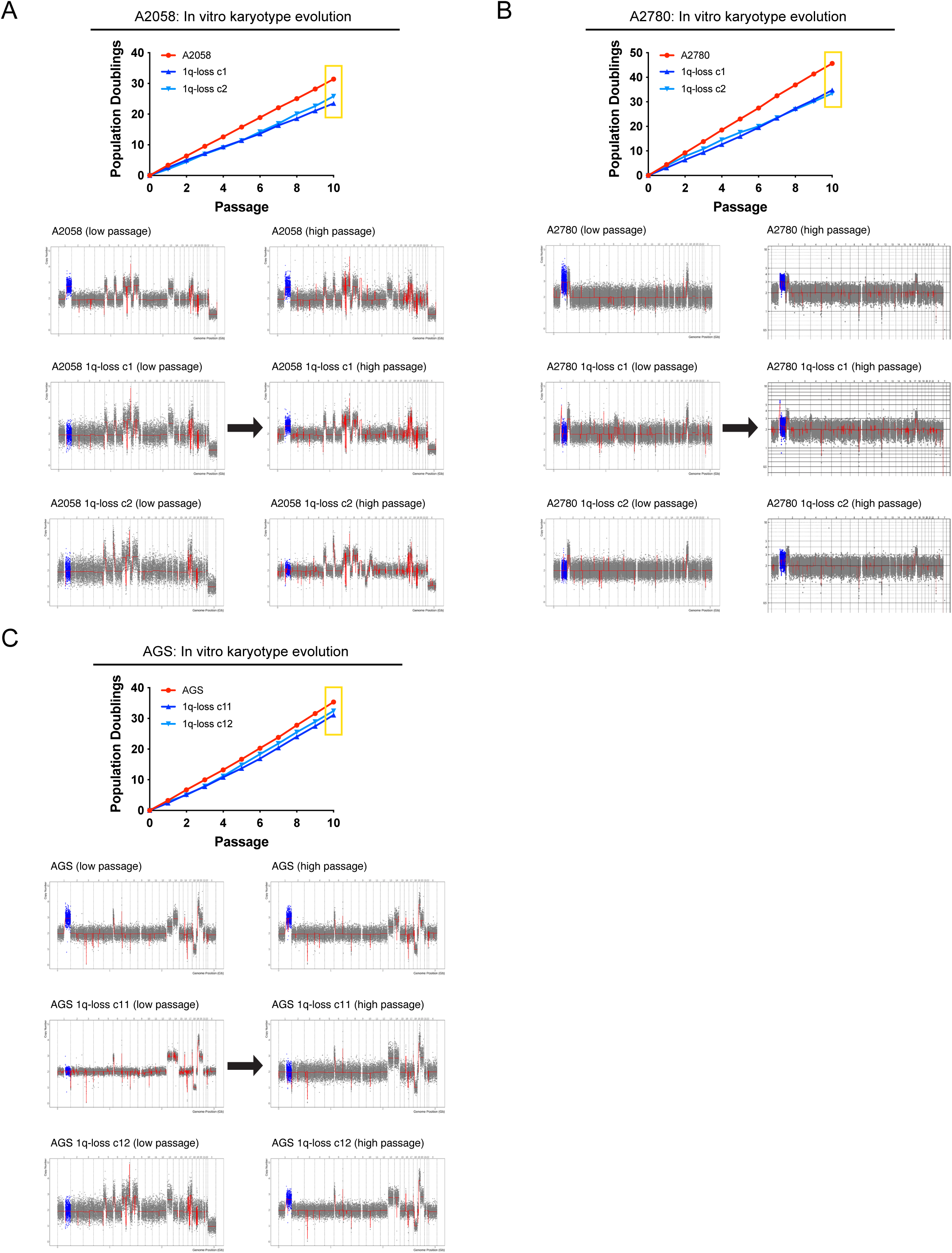
Karyotype evolution of 1q-disomic cancer cells following serial passaging *in vitro*. (A) Regain of the 1q-trisomy during prolonged *in vitro* passaging of A2058 1q-disomic clones. (B) Regain of the 1q-trisomy during prolonged *in vitro* passaging of A2780 1q-disomic clones. (C) Regain of the 1q-trisomy during prolonged *in vitro* passaging of AGS 1q-disomic clones. For A-C, cells were harvested at the start of the proliferation assay and after 10 passages at the conclusion of the assay, and subject to SMASH karyotyping. Karyotypes are shown below, with chromosome 1q highlighted in blue.

**Figure S13.**
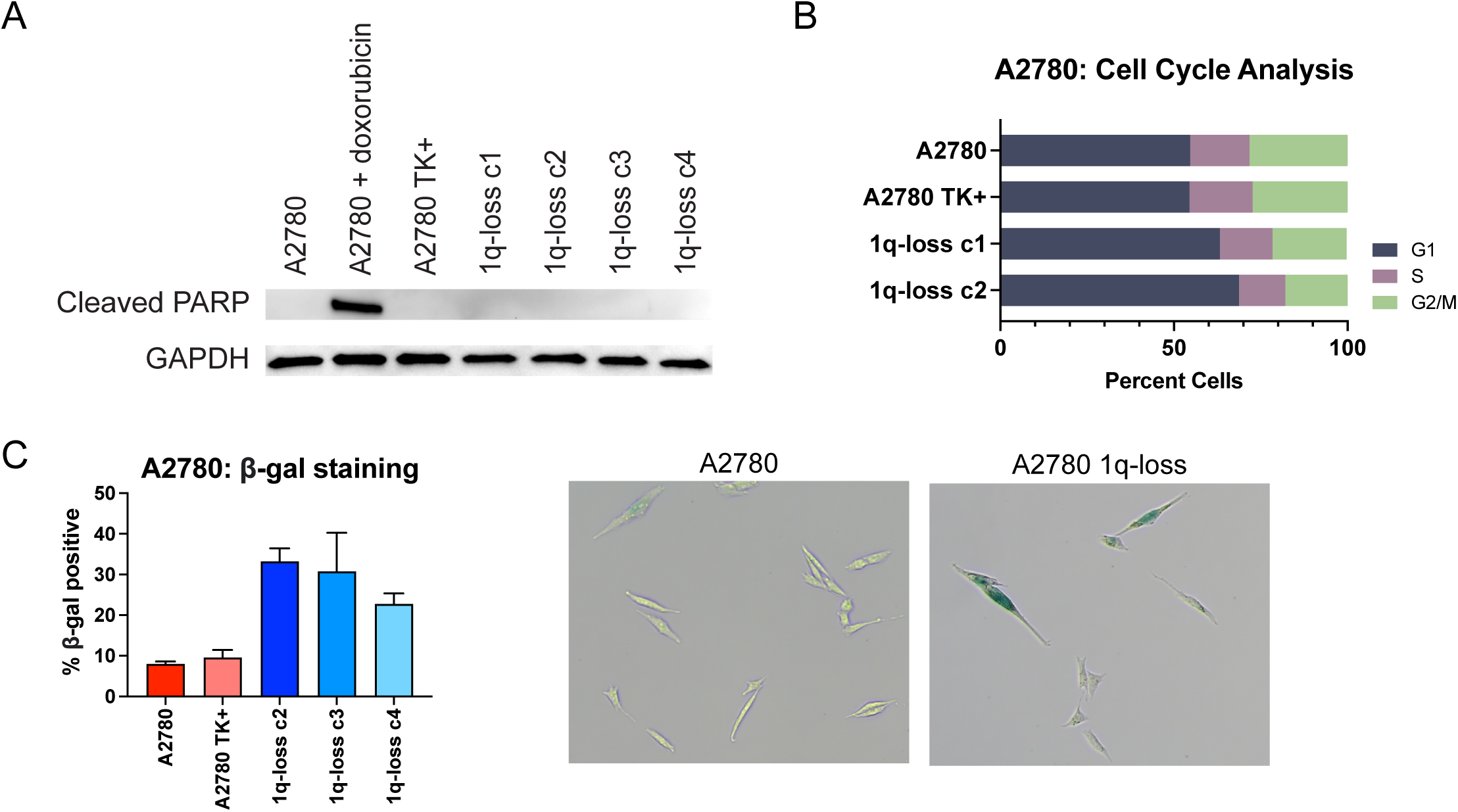
Elimination of the 1q trisomy causes a G1 delay and an increase in senescence. (A) Levels of cleaved PARP1 were assessed using western blotting to determine whether aneuploidy-loss induces apoptosis. As a positive control, wild-type cells were treated with the DNA-damaging agent doxorubicin. GAPDH was analyzed as a loading control. (B) Cell cycle analysis of A2780 and A2780 1q-disomic clones, determined via propidium iodide staining. (C) A2780 and A2780 1q-disomic clones were stained for beta-galactosidase expression to assess the levels of senescence in each population.

**Figure S14.**
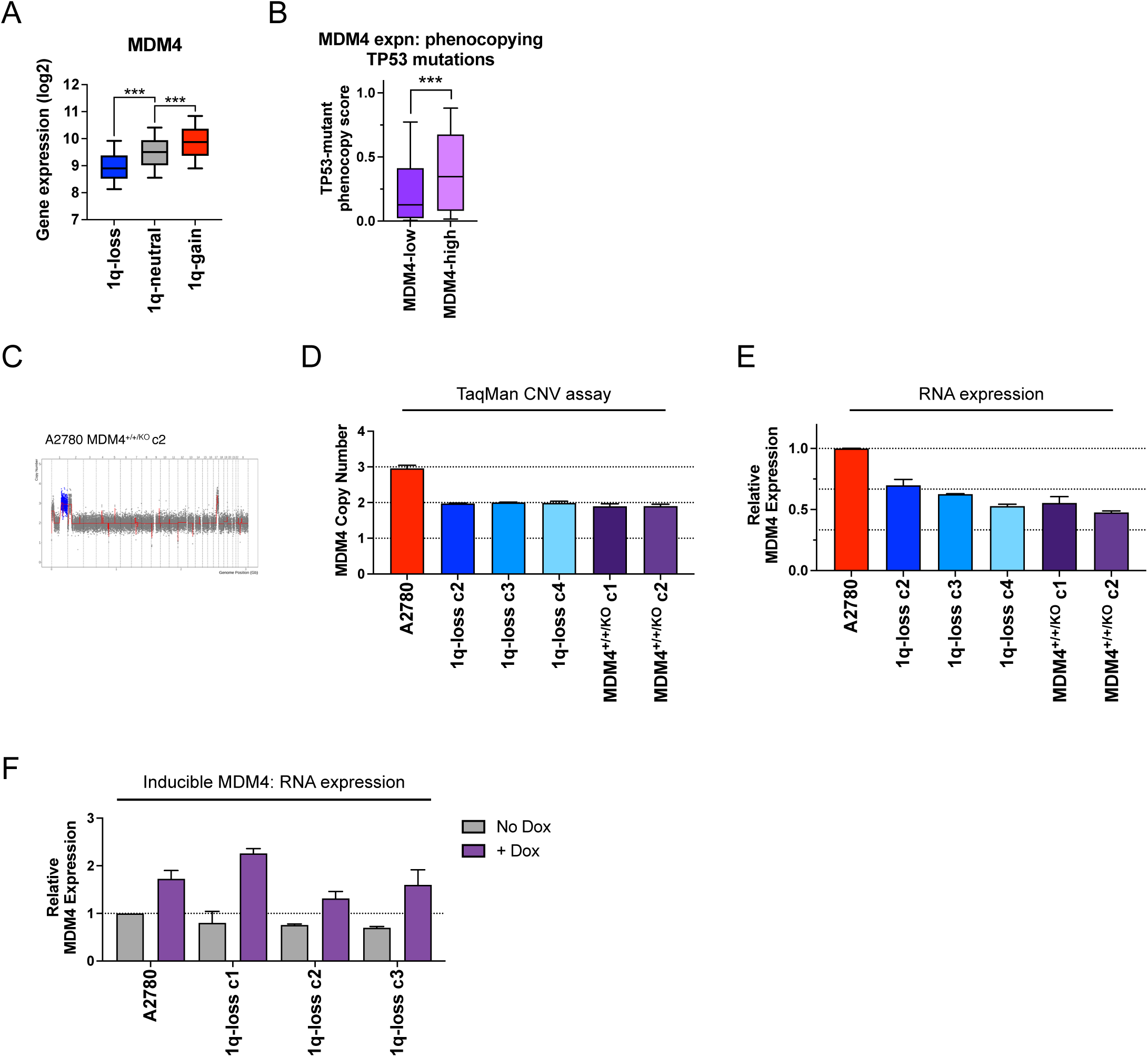
MDM4 expression in human tumors and MDM4 genetic manipulations in A2780 cells. (A) Boxplots displaying the expression of MDM4 in human cancers from TCGA split based on the copy number of chromosome 1q. (B) Boxplots displaying the TP53-mutation phenocopy signature^57^ in cancers from the TCGA, split based on the expression of MDM4. Only TP53-wildtype cancers are included in this analysis. (C) SMASH karyotype demonstrating maintenance of the chromosome 1q trisomy in an MDM4^+/+/KO^ clone. Chromosome 1q is highlighted in blue. (D) TaqMan copy number verification of the deletion of a single copy of MDM4 in A2780 cells. Mean ± SEM, n=2 probes for MDM4. (E) MDM4^+/+/KO^ and 1q-disomic clones exhibit decreased expression of MDM4, as determined through TaqMan gene expression assays. Data are normalized to parental A2780. (F) Expression of MDM4 in 1q-trisomic and 1q-disomic cells harboring MDM4 under the control of a doxycycline-inducible promoter. MDM4 expression was measured using qRT-PCR. ***p < 0.0005

**Figure S15.**
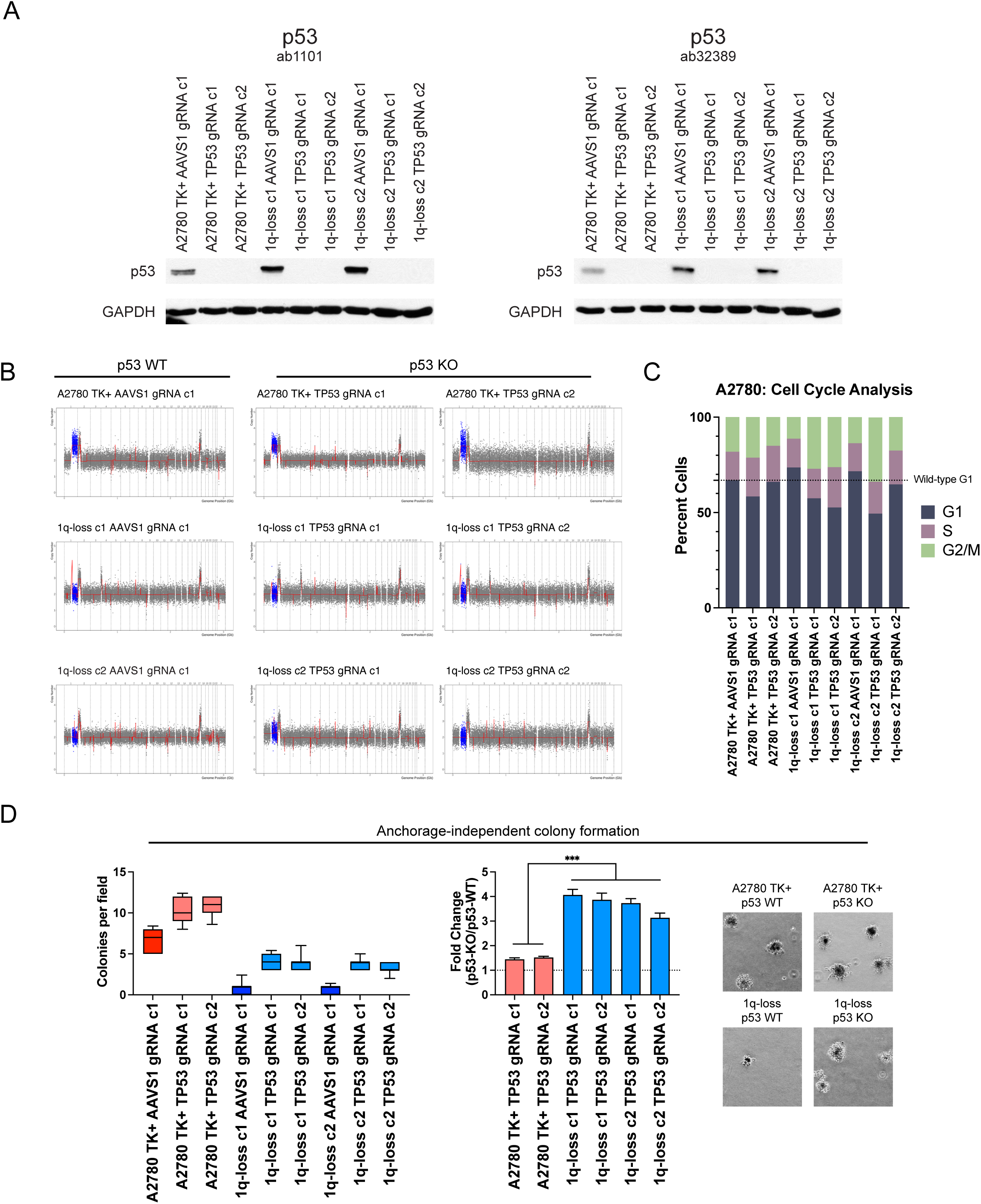
Deletion of TP53 selectively increases the fitness of 1q-disomic cells. (A) Western blot validation of p53-KO with two different antibodies. AAVS1 gRNA clones serve as isogenic p53- WT controls. GAPDH was used as a loading control. (B) SMASH karyotypes of p53-KO and p53-WT clones. TK+ integrant clones maintain the 1q-trisomy, 1q-disomic clones maintain the 1q-disomy, and no other karyotypic alterations are observed. Chromosome 1q is highlighted in blue. (C) Cell cycle analysis of p53-KO and p53-WT clones in A2780 TK+ and 1q-disomic lines, determined via propidium iodide staining. (D) Deletion of TP53 increases anchorage independent growth by ∼1.5-fold in 1q-trisomic clones and by ∼4-fold in 1q-disomic clones. Boxes represent the 25th, 50th, and 75th percentiles of colonies per field, while the whiskers represent the 10th and 90th percentiles. Unpaired t-test, n = 15 fields of view, data from representative trial. Representative images are shown. ***p < 0.0005

**Figure S16.**
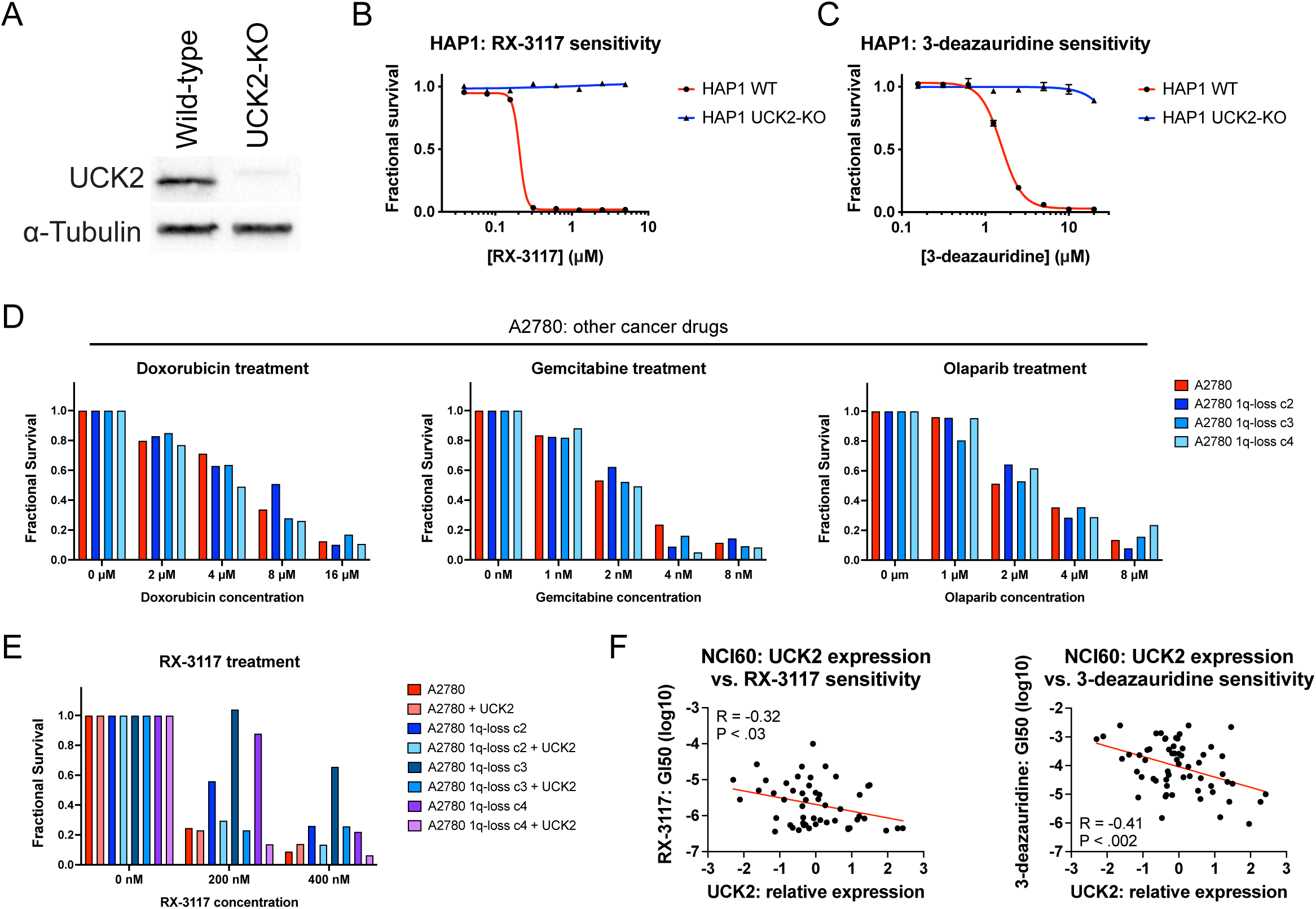
UCK2 expression sensitizes cancer cells to RX-3117 and 3-deazauridine. (A) Western blotting verifying lack of UCK2 expression in the HAP1 UCK2-KO clone. Tubulin levels were analyzed as a loading control. (B) 7-point dose response curve displaying the viability of HAP1 and HAP1 UCK2-KO cells treated with varying concentrations of RX-3117. (C) 7-point dose response curve displaying the viability of HAP1 and HAP1 UCK2-KO cells treated with varying concentrations of 3-deazauridine. (D) Viability of A2780 cells treated with different concentrations of the indicated anti-cancer drug. (E) Viability of A2780 cells or A2780 cells transduced with UCK2 cDNA treated with RX-3117. (F) Correlation between UCK2 expression and sensitivity to RX-3117 (left) or 3-deazauridine (right) across the NCI-60 panel of cancer cell lines^69^.

